# OPA1-dependent mitochondrial remodeling coordinates TCR signaling and metabolic adaptation during iNKT cell differentiation

**DOI:** 10.64898/2026.09.12.751163

**Authors:** Sophia P. M. Sok, Jianan Cheng, Rufei Lu, Tommy L Lewis, Hiromi Sesaki, Renata O Pereira, Kai Sun, Victoria Sun, Meng Zhao

**Author notes:** Corresponding authors: Meng Zhao, Ph.D. These authors contributed equally (Sophia P. M. Sok, Jianan Cheng). Disclosures: The authors declare no competing financial interests.

## Abstract

Invariant natural killer T (iNKT) cells require mitochondrial metabolism for terminal effector development. We found iNKT cells express elevated levels of proteins regulating mitochondrial membrane dynamics, and identified Opa1, but not Drp1, as selectively required for iNKT cell differentiation. OPA1 deficiency disrupted mitochondrial cristae organization, reduced mitochondrial membrane potential, increased mitochondrial mass, and altered calcium homeostasis in iNKT cells. Bulk and single-cell transcriptomic analyses revealed impaired TCR-responsive gene expression, activation of mitochondrial stress adaptation and integrated stress response, enhanced glycolysis, and retention of immature differentiation features. SCENITH analysis demonstrated increased glycolytic dependence; LDHA became required in Opa1-deficient iNKT cells while dispensable for normal iNKT development, indicating compensatory glycolytic adaptation. EGTA in thymic organ culture partially restored NKT1 marker expression in Opa1-deficient cells. Co-deletion of Drp1 improved mitochondrial morphology, TCR signaling, stress and metabolic adaptation, and partially rescued iNKT cell differentiation, demonstrating that balanced mitochondrial dynamics coordinate mitochondrial function and terminal effector development.

## Introduction

Invariant natural killer T (iNKT) cells undergo agonist selection in the thymus and follow a developmental program directed by a distinct combination of TCR, SLAM family, and cytokines^1^. While positively selected conventional thymocytes undergo lineage commitment, tolerance induction and maturation into quiescent naïve T cells, iNKT cell precursors instead undergo a c-Myc-dependent proliferative burst while concurrently differentiating into NKT1, NKT2 and NKT17 effector subsets^2^. This unique developmental trajectory may impose exceptional metabolic demands. Accordingly, c-Myc is required for post-selection transition^3, 4^, whereas mTORC1 and mTORC2 regulate complementary aspects of proliferation, maturation and subset specification during iNKT cell development ^5–9^. Among the metabolic pathways supporting iNKT cell development, mitochondria are uniquely positioned to integrate bioenergetics with intracellular signaling. Activated iNKT cells preferentially channel glucose into the pentose phosphate pathway and mitochondrial oxidative metabolism rather than lactate production and depend on oxidative phosphorylation for optimal survival and effector function^10^. Genetic disruption of Rieske iron sulfur protein (RISP), a component of the complex III of the mitochondrial electron transport chain (ETC), demonstrated that intact mitochondrial respiration is required for iNKT cell development and TCR signaling^11^. In parallel, we showed that thymic iNKT cells possess greater mitochondrial content and activity than conventional T cells, and that mitochondrial calcium buffering tunes TCR signaling to direct iNKT cell differentiation^12, 13^. Together, these studies establish mitochondria as both metabolic and signaling hubs that are fundamental to iNKT cell development and function.

Mitochondria are highly dynamic organelles whose morphology, ultrastructure, and intracellular distribution are continuously remodeled through coordinated fusion, fission, and intracellular trafficking^14–16^. These processes regulate cristae structure, respiratory complex organization, calcium homeostasis, and metabolic adaptation, thereby influencing cellular signaling and fate decisions^17–19^. In conventional T cells, early deletion of the inner membrane fusion protein OPA1 arrests thymocyte maturation at the DN3/β-selection checkpoint, whereas deletion of the fission protein DRP1 reduces thymocyte cellularity without altering subset representation^20, 21^. Beyond thymic development, in mature CD8 T cells, OPA1 was initially shown to be selectively required for memory T cell formation through cristae dependent metabolic programming, whereas a recent study instead identified a prominent requirement for OPA1 during effector T cell proliferation and survival, including functions beyond mitochondrial fusion^22, 23^. OPA1 is separately required for Th17 effector function, where it couples TCA cycle metabolism to LKB1-dependent transcriptional remodeling^24^. Whether mitochondrial dynamics similarly orchestrate the unique metabolic and signaling requirements underlying agonist selected iNKT cell differentiation remains unknown.

Here we show that thymic iNKT cells express high levels of mitochondrial membrane remodeling proteins while maintaining a more punctate mitochondrial network than conventional CD4 T cells. Conditional deletion of the inner membrane fusion protein OPA1, but not the fission protein DRP1, skewed iNKT cell differentiation toward immature precursor and NKT2-like states while impairing terminal NKT1 and NKT17 cell differentiation. OPA1 deficiency disrupted mitochondrial structure and function, calcium homeostasis, TCR signaling, and metabolic adaptation. Calcium chelation during thymic organ culture partially corrected the differentiation defect of OPA1- deficient iNKT cells, supporting a functional role for calcium dysregulation in impaired NKT1 development.

OPA1-deficient cells acquired a genetic dependence on LDHA, demonstrating that compensatory glycolytic rewiring becomes functionally required. Single-cell transcriptomic analysis revealed that OPA1-deficient NKT1 cells failed to acquire the canonical NKT1 program and instead adopted an alternative state marked by mitochondrial stress and integrated stress response signatures. Co-deletion of DRP1 broadly ameliorated the mitochondrial, metabolic, signaling, and transcriptional abnormalities caused by OPA1 deficiency and normalized iNKT subset distribution, identifying balanced mitochondrial membrane dynamics as a key regulator of iNKT cell differentiation.

## Results

### Thymic iNKT cells possess a distinct mitochondrial shape and require OPA1 for development

To determine whether thymic iNKT cells possess specialized mitochondrial organization, we imaged mitochondria in freshly sorted iNKT cells and conventional CD4⁺ thymocytes from mito-Dendra2 reporter mice^25^. We previously showed, using this reporter, that iNKT cells contain greater mitochondrial volume than conventional T cells^12^. Here, mitochondria in CD4⁺ T cells formed elongated tubular structures, whereas iNKT cells displayed a more compact and punctate mitochondrial pattern, consistent with a distinct organization of their expanded mitochondrial compartment (Fig. 1A). Next, we performed quantitative proteomic analysis of freshly isolated thymic iNKT and CD4⁺ T cells. Relative to CD4⁺ thymocytes, iNKT cells showed increased abundance of proteins involved in mitochondrial membrane remodeling and organization, including the core fusion and fission GTPases OPA1^26, 27^ and DRP1^28^, the outer-membrane fusion-fission site factor SLC25A46^29^, the inner-membrane/cristae-associated proteins TMEM11^30^ and GHITM^31^, and mitochondrial trafficking regulators RHOT1, RHOT2, and ARMC10^32, 33^. In contrast, components of the MICOS-SAM (MIB) network were reduced, including the MICOS subunits APOO/MIC26 and APOOL/MIC27^34^ and the outer membrane SAM subunit MTX1^35^ (Fig. 1B). Together with their distinct mitochondrial morphology, this proteomic profile suggested that mitochondrial membrane remodeling may be particularly important in thymic iNKT cells.

**Figure 1.**
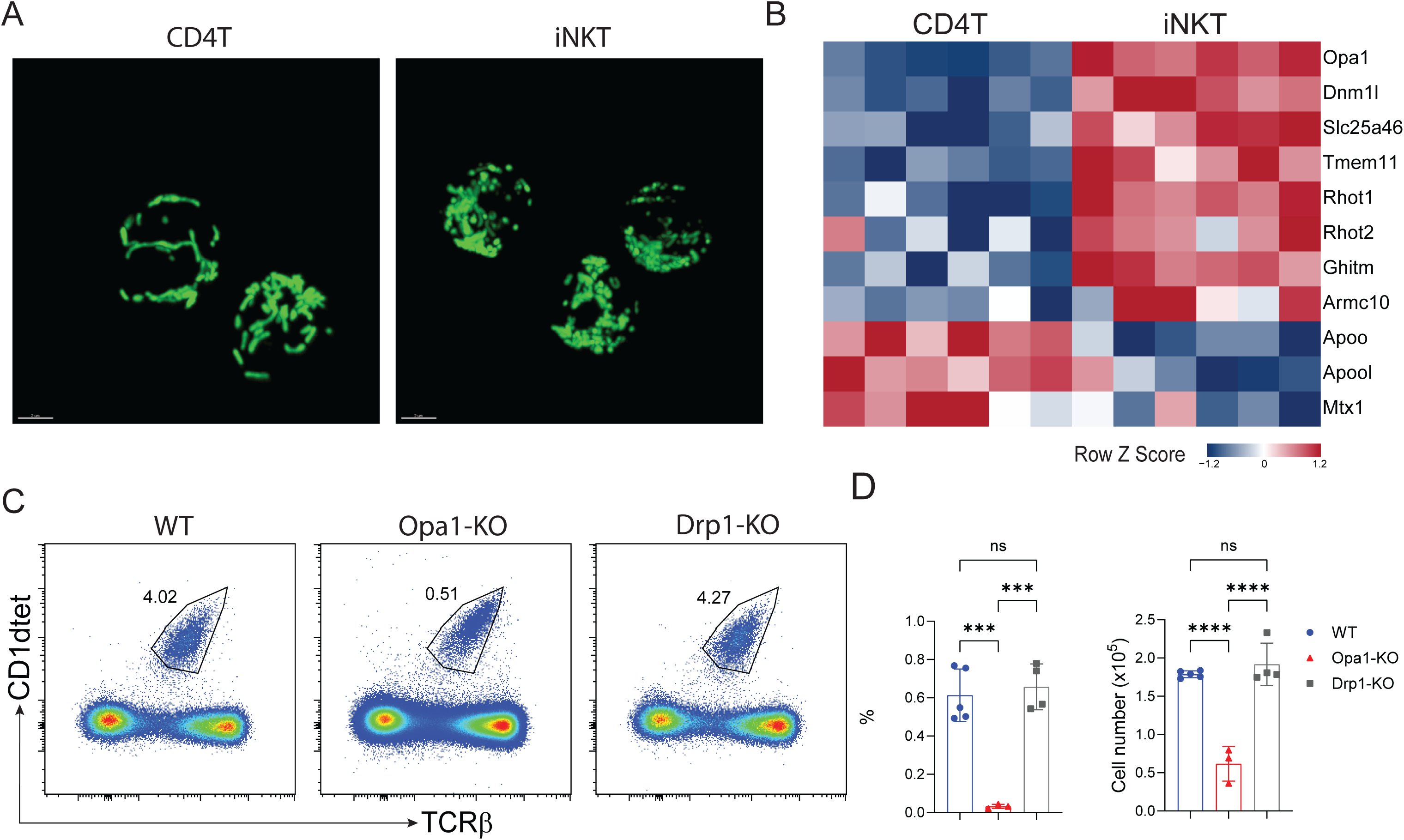
Thymic iNKT cells display mitochondrial architecture distinct from CD4 T cells and selectively require Opa1 for development. (A) Representative confocal images of mitochondria in freshly isolated thymic CD4^+^ T cells and iNKT cells from mitoDendra2 reporter mice. (B) Heatmap showing relative abundance of representative differentially expressed mitochondrial membrane dynamics proteins in naïve thymic CD4^+^ T cells and iNKT cells determined by quantitative proteomics. Columns represent biological replicates. (C) Representative flow cytometry plots showing thymic iNKT cells within CD8^-^ thymocytes from WT, Opa1-KO and Drp1-KO mice. (D) Frequency within total live cells (left) and cell number (right) of total thymic iNKT cells in WT, Opa1-KO and Drp1-KO mice. Bars indicate mean ± SD. Data are representative results from three independent experiments. Statistical significance was determined by one-way ANOVA with Turkey’s multiple comparisons test. test. ns, not significant; ***P < 0.001; ****P < 0.0001.

To test the functional requirements for the core fusion and fission machinery, we crossed Opa1-fl/fl and Dnm1l/Drp1-fl/fl mice with CD4-Cre mice to delete Opa1 or Drp1 at CD4+CD8+ double positive stage. Consistent with a previous report ^36^, Opa1 deficiency markedly reduced thymic iNKT cell frequency and absolute number (Fig. 1C,D). Surprisingly, despite more punctate mitochondrial morphology in iNKT cells, Drp1 deficient mice showed iNKT cell frequencies and numbers comparable to those of littermate WT controls. Conventional αβ T cell development remained largely intact in both mutant strains. Thus Opa1, but not Drp1, is selectively required for thymic iNKT cell development.

### OPA1 cell intrinsically promotes terminal iNKT cell differentiation

Opa1-KO mice exhibited a marked reduction in mature S3 cells with accumulation of S2 cells, whereas S0 and S1 populations were largely preserved, indicating that OPA1 is dispensable for positive selection and early precursor development but required during terminal maturation (Supplemental Fig. S1A-C). Analysis of effector subsets likewise revealed a pronounced reduction in NKT1 cells accompanied by reciprocal expansion of NKT2 cells, while the modest reduction in NKT17 cells did not remain significant after correction for multiple comparisons (Supplemental Fig. S1D-F).

To distinguish cell intrinsic from extrinsic effects, congenically marked WT (CD45.1) bone marrow was mixed 1:1 with either WT or OPA1-KO (CD45.2) bone marrow and transplanted into lethally irradiated recipients. Within the same thymic environment, OPA1-KO donor cells generated substantially fewer total iNKT cells than their paired WT counterparts, despite largely normal conventional αβ T cell development (Supplemental Fig. S1G-I). NKT1 cell reduction and NKT2 cell expansion closely mirrored the phenotype observed in intact mice (Supplemental Fig. S1J-L), demonstrating that these differentiation defects are cell intrinsic. To exclude defects upstream of terminal differentiation, we examined the DP thymocyte compartment and CD1d expression on selecting thymocytes, both of which were comparable between genotypes (Supplemental Fig. S1M-P). Likewise, enforced expression of the anti-apoptotic protein BCL-XL failed to restore total iNKT cell numbers or normalize subset distribution (Supplemental Fig. S1Q-T), indicating that impaired differentiation cannot be explained by defective selection or BCL-XL-sensitive apoptosis. Conversely, deletion of Opa1 using Tbet-Cre^37^ substantially reduced OPA1 expression in NKT1 cells but did not alter total iNKT cellularity or subset distribution (Supplemental Fig. S2). Together, these findings define a developmental window in which OPA1 acts cell intrinsically after iNKT cell selection but before acquisition of the mature NKT1 state.

### Genetic inhibition of mitochondrial fission factor DRP1 reverses iNKT cell developmental defects caused by OPA1 deficiency

OPA1 promotes mitochondrial inner membrane fusion, whereas DRP1 mediates mitochondrial fission. We therefore hypothesized that Opa1 deficiency causes unopposed DRP1-dependent mitochondrial fragmentation. To test this, we generated Opa1-f/f; Drp1-f/f; CD4cre (DKO) mice. Deletion of Drp1 alone did not significantly alter iNKT cell frequency, number, or subset distribution (Fig. 1). In contrast, concomitant deletion of Drp1 substantially recovered thymic iNKT cell frequency and number in Opa1-KO mice, although neither returned fully to WT levels (Fig. 2A-C). Thus, the iNKT cell developmental defect caused by Opa1 loss requires DRP1, a genetic interaction consistent with excessive mitochondrial fission.

**Figure 2.**
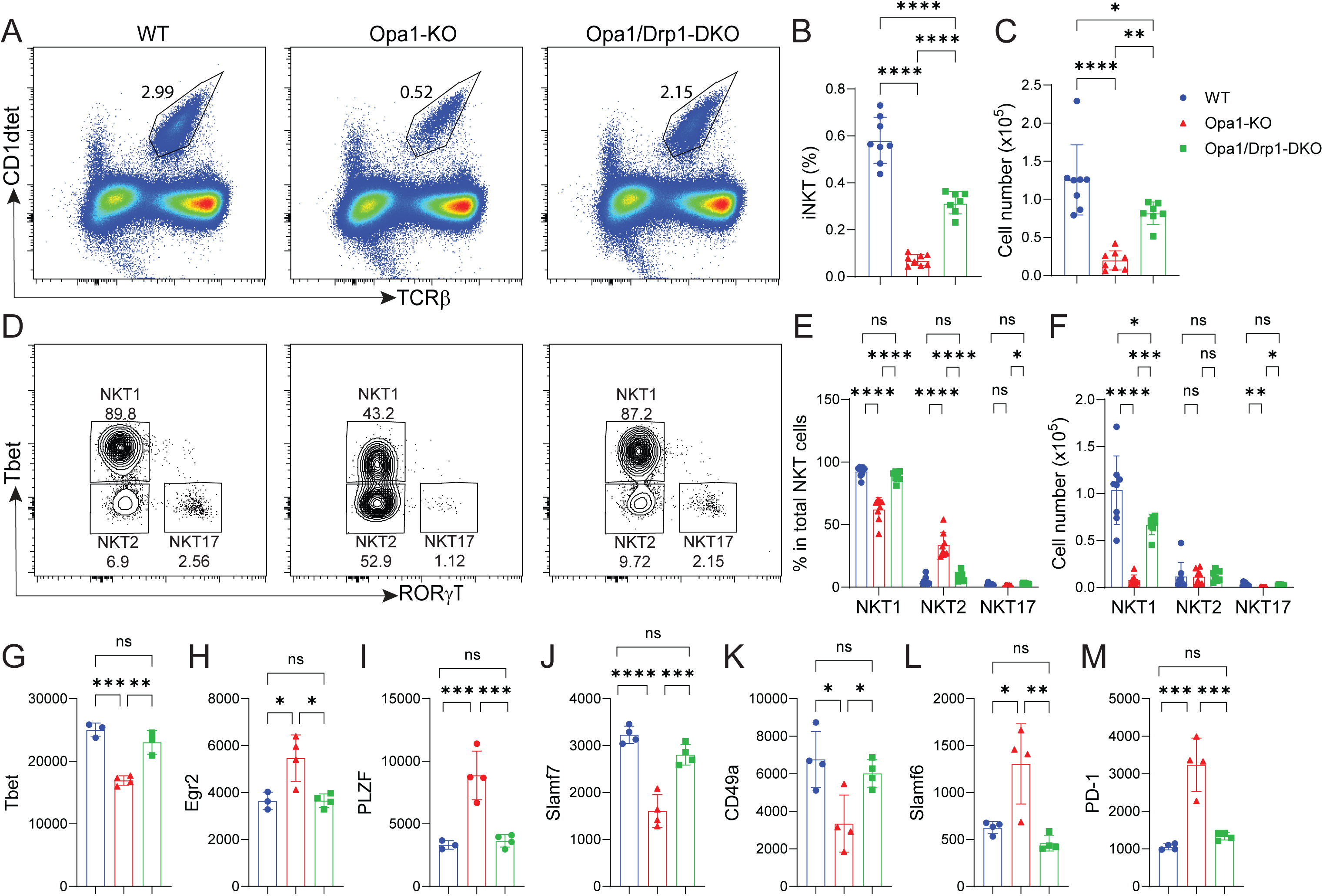
Drp1 co-deletion restores iNKT cell subset specification and partially restores thymic iNKT cellularity in Opa1 deficient mice. (A) Representative flow cytometry plots showing total iNKT cells within CD8α^-^ thymocytes, (B) frequency within total live cells, and (C) cell number of thymic iNKT cells from WT, Opa1-KO and Opa1/Drp1-DKO mice. (D) Representative flow cytometry plots showing NKT1 (TBET^+^RORγt^-^), NKT2 (TBET^-^RORγt^-^) and NKT17 (TBET^-^ RORγt^+^) subsets. (E) Frequency within total iNKT cells and (F) absolute numbers of thymic iNKT cell subsets. (G-M) Expression of transcription factors and surface markers in thymic NKT1 cells. (B, C, E, F) are pooled from three independent experiments. (G-M) are representative results from three independent experiments. Bars indicate mean ± SD. Statistical significance was determined by one-way ANOVA with Tukey’s multiple comparisons test, performed separately within each subset in (E) and (F). ns, not significant; *P < 0.05; **P < 0.01; ***P < 0.001; ****P < 0.0001.

Furthermore, Drp1 deletion fully normalized the distribution of iNKT cell subsets (Fig. 2D, E). Absolute NKT1 numbers significantly increased in DKO relative to Opa1-KO mice but remained below WT, mirroring the partial rescue of total iNKT cell cellularity (Fig. 2F). We next asked whether the residual NKT1 cells in Opa1-KO mice acquired a mature NKT1 phenotype. Opa1-KO NKT1 cells expressed lower TBET together with elevated levels of the precursor/NKT2-associated transcription factor EGR2 and PLZF (Fig. 2G-I). Consistently, the mature NKT1 surface markers SLAMF7 and CD49A were reduced, while NKT2 associated markers SLAMF6 and PD- 1 were increased in Opa1-KO NKT1 cells (Fig. 2J-M). Strikingly, the phenotypic identity of NKT1 cells were fully normalized by Drp1 co-deletion. Together, these findings suggest that both the block in terminal NKT1 differentiation and the immature phenotype of the residual NKT1 cells in Opa1-deficient mice arise largely from disruption of the normal balance between mitochondrial fusion and fission.

### Balanced mitochondrial dynamics maintain mitochondrial architecture and function

To define how Opa1 loss alters mitochondrial organization in iNKT cells, we first examined mitochondrial morphology in freshly isolated cells from mito-Dendra2 reporter mice. WT NKT1 cells contained a heterogeneous mitochondrial network composed of both punctate and short tubular structures, whereas Opa1- KO NKT1 cells were dominated by small punctate mitochondria, consistent with impaired mitochondrial fusion (Fig. 3A). Co-deletion of Drp1 largely restored the mixed punctate/tubular morphology (Fig. 3A). Quantification of the Dendra2 signal further showed that total mitochondrial volume was increased in Opa1-KO cells and normalized by Drp1 co-deletion (Fig. 3B), indicating that Drp1 co-deletion normalizes mitochondrial morphology and content.

**Figure 3.**
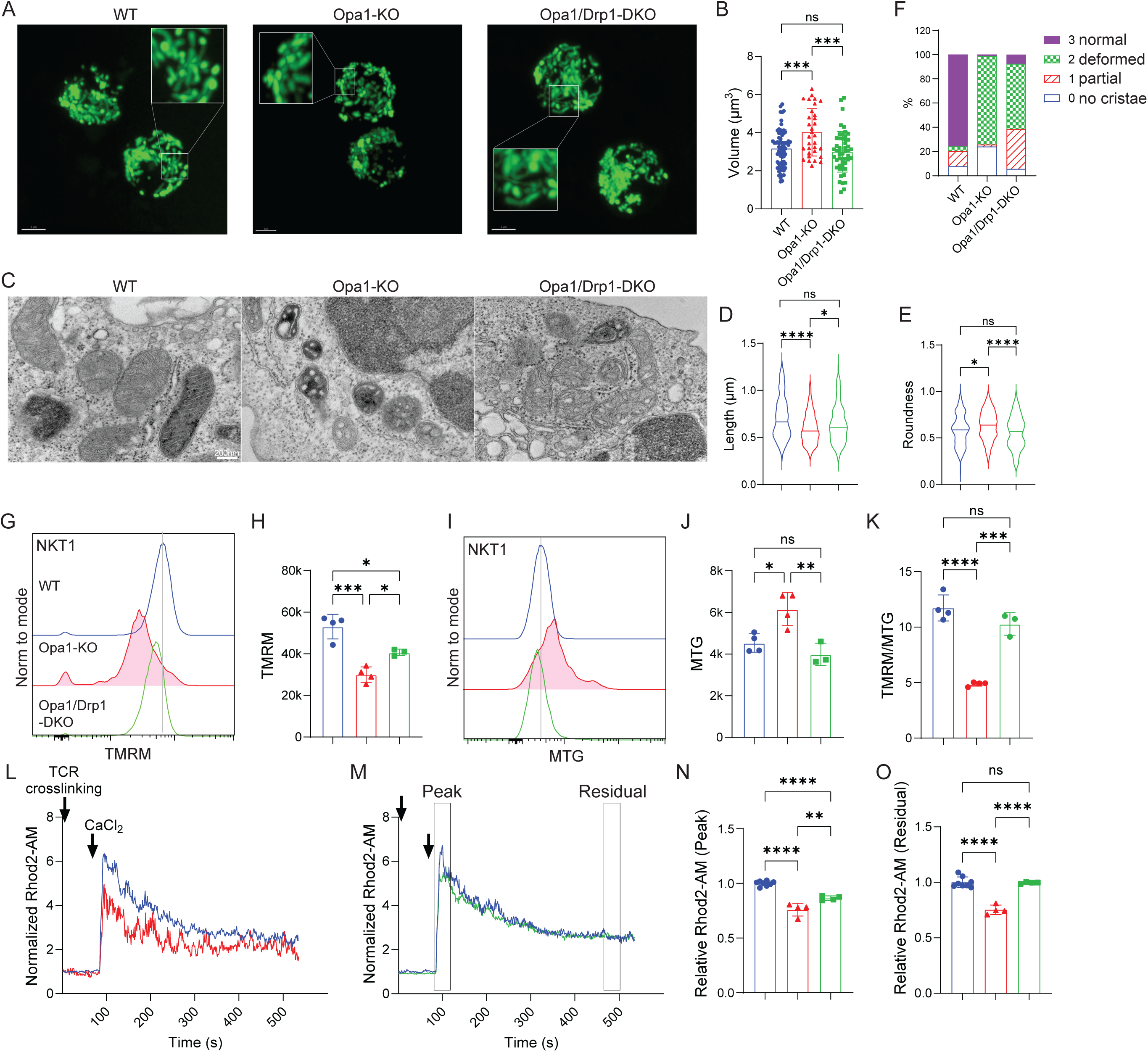
Drp1 co-deletion improves mitochondrial morphology and function in Opa1 deficient iNKT cells. (A) Representative confocal images of mitochondria in freshly isolated thymic NKT1 cells from mito-Dendra2 reporter mice. (B) Quantification of mitochondrial volume per cell based on mito-Dendra2 images. (C) Representative transmission electron micrographs of thymic NKT1 cell (ICOS^lo^) mitochondria. (D-F) Quantification of mitochondrial length (D), roundness (E), and blinded cristae integrity score (F). (G-K) Mitochondrial membrane potential and mitochondrial mass in thymic NKT1 cells. Representative histograms of TMRM (G) and MitoTracker Green (MTG; I) staining and quantification of TMRM fluorescence (H), MTG fluorescence (J), and the TMRM/MTG ratio (K). (L-O) Mitochondrial calcium level following TCR stimulation. Representative traces comparing WT with Opa1-KO (L) or with Opa1/Drp1-DKO (M) cells. Boxes in (M) indicate the intervals used to quantify peak and residual Rhod2-AM signals. Quantification of peak (N) and residual (O) Rhod2-AM signal, each normalized to the pre-stimulation baseline of the same trace and expressed relative to the WT mean within each experiment. Each point in (B) represents an individual cell pooled from three independent experiments. (D-F) summarize individual mitochondria quantified from transmission electron micrographs obtained in one representative experiment. (G-O) show representative results from three independent experiments. In (B, H, J, K, N, O), bars indicate mean ± SD and symbols represent individual cells (B), individual mice (H, J, K) or individual time points (N, O). In (D, E), violin plots show the median and quartiles. Statistical significance was determined by one-way ANOVA with Tukey’s multiple comparisons test. ns, not significant; *P < 0.05; **P < 0.01; ***P < 0.001; ****P < 0.0001.

To further characterize mitochondrial ultrastructure, we performed transmission electron microscopy of freshly isolated thymic iNKT cells. Consistent with the Dendra2 imaging, Opa1-KO mitochondria appeared shorter and more circular than WT mitochondria, and both were restored by Drp1 co-deletion (Fig. 3C-E). In contrast, while WT mitochondria predominantly contained intact lamellar cristae, Opa1-KO mitochondria frequently exhibited severely disrupted or absent cristae, and normal cristae architecture was not fully restored in DKO (Fig. 3C, F). Thus, Drp1 deletion efficiently rescues mitochondrial morphology despite persistent defects in OPA1- dependent inner membrane organization.

We next asked whether restoration of mitochondrial morphology improved mitochondrial function. As iNKT cells express high levels of the multidrug transporter MDR1/P-glycoprotein, which actively extrudes both TMRM and MitoTracker dyes, all staining was performed in the presence of the MDR1 inhibitor PSC833 to eliminate dye efflux artifacts^12^. Opa1-KO iNKT cells exhibited significantly reduced TMRM fluorescence compared with WT cells, whereas Drp1 deletion substantially improved it (Fig. 3G, H, Supplemental Fig. S3A, B, F, G). In contrast, staining with MitoTracker Green (MTG), which reflects mitochondrial mass independently of membrane potential, was increased on Opa1-KO NKT1 cells^36^ and restored in DKO cells (Fig. 3I, J, Supplemental Fig. S3C, D, H, I), consistent with findings from Dendra2 imaging (Fig. 3A, B). Consequently, the ratio of TMRM to MTG, an indicator of membrane potential normalized to mitochondrial mass, was markedly reduced by Opa1 deficiency and restored in Opa1/Drp1-DKO (Fig. 3K, Supplemental Fig. S3E, J). In contrast, conventional CD4 T cells exhibited no detectable abnormalities, whereas CD8 T cells showed only a modest reduction in TMRM/MTG ratio (Supplemental Fig. S3K-T). Together, Opa1 deficiency leads to the accumulation of an expanded but functionally compromised mitochondrial network selectively within iNKT cells, a defect that is substantially corrected by co-deletion of Drp1.

Because mitochondrial calcium uptake depends on mitochondrial membrane potential^38^, we next examined mitochondrial calcium dynamics following TCR stimulation. WT and mutant thymocytes were differentially labeled with CFSE, mixed within the same sample, loaded with the mitochondria targeted calcium indicator Rhod2-AM^39^, and analyzed simultaneously to ensure identical dye loading, stimulation, and acquisition conditions. Following TCR crosslinking and extracellular CaCl₂ addition, Opa1-KO NKT1 cells exhibited significantly reduced mitochondrial calcium uptake, with both the initial peak and sustained post-stimulation calcium signal diminished relative to paired WT cells (Fig. 3L-O). Reversal of CFSE labeling produced identical results (Supplemental Fig. S3U-X). Drp1 co-deletion fully restored the sustained mitochondrial calcium signal but only partially rescued the initial peak response (Fig. 3L-O). Together, these findings indicate that restoration of mitochondrial morphology is accompanied by substantial recovery of mitochondrial membrane potential, although OPA1-dependent features, including cristae organization and rapid mitochondrial calcium uptake, remain incompletely recovered.

### OPA1 deficiency reshapes the thymic iNKT cell developmental landscape and reveals a distinct NKT1 transcriptional state

We next examined how these mitochondrial defects influence the developmental and transcriptional landscape of thymic iNKT cells. After quality filtering and doublet removal, 21,512 high-quality iNKT cells were retained for downstream single cell RNAseq (scRNAseq) analyses (Supplemental Fig. S4). Detected genes, UMI counts, and mitochondrial transcript percentages, were comparable across genotypes (Supplemental Fig. S4A-C). Unsupervised clustering identified ten transcriptionally distinct clusters (Fig. 4A). Clusters were annotated according to established developmental and lineage markers as Stage 0, NKT precursor (NKTp), NKT2, NKT17, and four transcriptionally distinct NKT1 populations (Supplemental Figs. S5). Stage 0 cells expressed Cd24a, Egr2, and Sox4, whereas proliferating precursor clusters were enriched for Mki67 and Top2a (Supplemental Fig. S5A). NKT2 cluster exhibited increased expression of Il4, Ccr4, and Il17rb, consistent with their known transcriptional overlap with precursor populations. NKT17 cells were defined by selective expression of Rorc, Il23r, Il1r1, and Ccr6, whereas NKT1 populations expressed the canonical effector program marked by Tbx21, Il2rb, Klrb1c, Nkg7, Ccl5, and Ifng (Supplemental Fig. S5A). Feature plots confirmed the expected spatial distribution of representative marker genes across the UMAP (Supplemental Fig. S5B). Detected gene and UMI counts varied across developmental clusters, with the highest transcript complexity observed in iNKT cell precursor populations and progressively lower complexity in differentiated NKT1 and NKT17 cells (Supplemental Fig. S4A-C)^40^. To benchmark these annotations against an independent reference, we reprocessed the raw scRNA-seq dataset from Krovi et al^40^. using the identical quality control workflow (Supplemental Fig. S6A). The regenerated UMAP reproduced the published results, allowing direct alignment of cluster identities between datasets (Supplemental Fig. S6A). Moreover, module scores derived from the Krovi cluster defining genes mapped to their corresponding clusters in our dataset, confirming strong concordance between the two atlases (Supplemental Fig. S6A, B).

**Figure 4.**
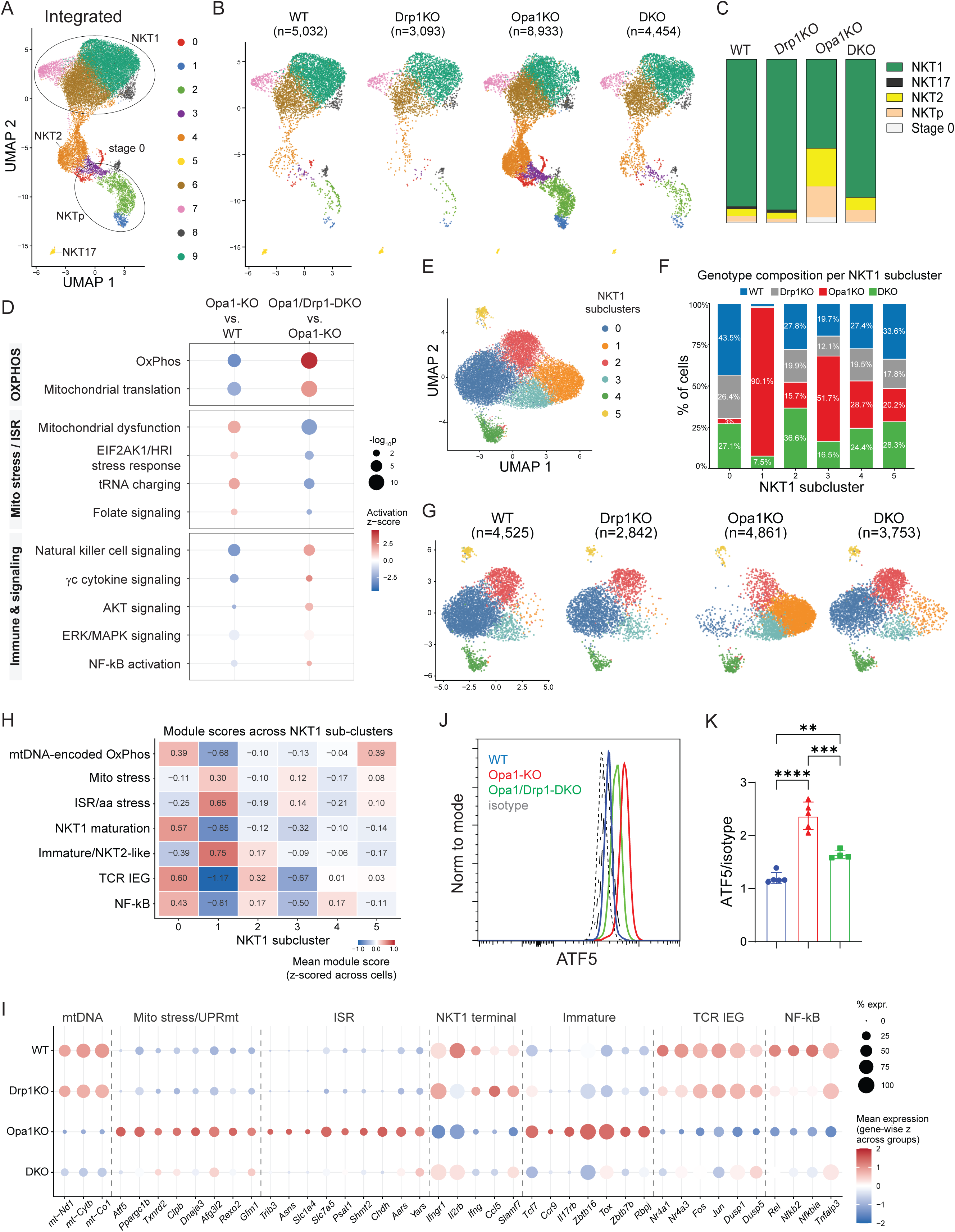
Opa1 deficiency redirects NKT1 cells into an alternative stress-associated transcriptional state that is reversed by Drp1 co-deletion. (A) UMAP of integrated thymic iNKT cells showing ten transcriptionally distinct clusters (C0-C9) corresponding to stage 0, NKTp, NKT2, NKT17, and NKT1 subsets. (B) Distribution of WT, Drp1-KO, Opa1-KO, and Opa1/Drp1-DKO cells on the integrated UMAP; n indicates cells recovered per genotype after quality filtering. (C) Subset composition per genotype, as the percentage of total iNKT cells. (D) Ingenuity Pathway Analysis of differentially expressed genes in NKT1 cells (Opa1-KO vs. WT and Opa1/Drp1-DKO vs. Opa1-KO). Dot color indicates activation z-score; dot size indicates −log₁₀(P). (E) UMAP of re-clustered NKT1 cells identifying six subclusters (SC0–SC5). (F) Genotype composition of each NKT1 subcluster. (G) Distribution of each genotype on the NKT1 subcluster UMAP. (H) Mean UCell module scores across NKT1 subclusters, z-scored across subclusters. (I) Representative genes from each module in (H). Dot color indicates mean expression z-scored across genotypes; dot size indicates the percentage of expressing cells. (J, K) Intracellular ATF5 expression in thymic NKT1 cells from WT, Opa1-KO, and Opa1/Drp1-DKO mice. (J) Representative flow cytometry histograms showing isotype control and ATF5 staining. (K) Quantification of ATF5 expression, shown as the ratio of ATF5 to isotype control MFI. Representative results from three independent experiments were shown. Bars represent mean ± SD. Statistical significance was determined by one-way ANOVA with Tukey’s multiple comparisons test. **P < 0.01; ***P < 0.001; ****P < 0.0001.

Visualization of individual genotypes on the integrated UMAP revealed that Opa1 deficiency selectively restructured the distribution of cells across the developmental continuum, whereas Drp1 deficiency alone closely resembled WT and largely restored the Opa1 deficient phenotype in DKO mice (Fig. 4B). Quantification of subset composition revealed expansion of Stage 0, NKTp, and NKT2 populations, accompanied by reductions in both NKT17 and NKT1 cells in Opa1-KO mice, consistent with flow cytometry results (Fig. 4C, Fig. 2). At higher resolution, cluster level analysis showed expansion of clusters 0-4 and a marked reduction of the terminal NKT1 cluster (cluster 9), all of which were largely restored by Drp1 co-deletion (Supplemental Fig. S6C). G2/M scores remained broadly comparable across genotypes (Supplemental Fig. S6D).

We next performed bulk RNA sequencing (RNAseq) of sorted thymic NKT1 and NKT2 cells from WT, Opa1-KO and Opa1/Drp1-DKO mice. Consistent with the established role of OPA1 in maintaining cristae structure and mitochondrial DNA integrity^17, 41^, Ingenuity Pathway Analysis (IPA) predicted suppression of oxidative phosphorylation and mitochondrial translation in Opa1-KO NKT1 cells (Fig. 4D, Supplemental Fig. S7A, B). In parallel, canonical pathways of mitochondrial integrated stress response, including EIF2AK1/HRI signaling, tRNA charging, and folate mediated one-carbon metabolism^42–44^, were predicted to be activated (Fig. 4D, Supplemental Fig. S7A, B). Consistent with the loss of mature NKT1 identity, pathways associated with NK cell function together with multiple downstream components of TCR and cytokine signaling, including γc cytokine, AKT, ERK/MAPK, and NF-κB signaling, were broadly suppressed (Fig. 4D, Supplemental Fig. S7A, B). Remarkably, most pathway alterations were substantially normalized by concomitant deletion of Drp1, indicating that the transcriptional consequences of Opa1 deficiency are largely driven by Drp1 dependent mitochondrial remodeling.

We next returned to the single cell dataset to determine how the bulk defined transcriptional programs were distributed within the heterogenous NKT1 compartment. NKT1 cells from all genotypes were re-clustered, recomputing variable features and dimensional reduction using this population alone, and resolving six subclusters, SC0-SC5 (Fig. 4E). WT and Drp1KO cells displayed highly similar subcluster distributions, whereas Opa1 deficiency markedly shifted cells toward SC1 and SC3 while depleting SC0, and these changes were substantially reversed by Drp1 co-deletion (Fig. 4G). Two additional subclusters (SC4 and SC5) were similarly represented across genotypes (Fig. 4G). Ucell module scores and expression levels of marker genes for a curated interferon stimulated gene (ISG) signature and a proliferation signature clearly distinguished these two subclusters (SC4 and SC5) respectively (Supplemental Fig. S8). The interferon responsive iNKT state (SC4) has been reported across organs and species^45^. The proliferative program in SC5 was substantially weaker than that observed in the dedicated cycling NKT precursor compartment (Supplemental Fig. S5), suggestive of a limited proliferative feature within NKT1 cells rather than contamination by the major precursor population.

To characterize the transcriptional programs distinguishing the NKT1 subclusters, we calculated UCell scores for seven curated gene modules representing mtDNA-encoded OXPHOS, mitochondrial maintenance and stress adaptation, ISR/amino acid stress, terminal NKT1 maturation, immature/NKT2-like retention, TCR-induced immediate early transcription, and NF-κB signaling, anchored in pathways and differentially expressed genes identified by the replicated bulk iNKT cell RNAseq analysis (Fig. 4D, Supplemental Fig. S7). Module scoring identified SC0 and SC1 as reciprocal transcriptional states (Fig. 4H). SC0 exhibited the strongest terminal NKT1 maturation, TCR immediate-early, NF-κB, and mtDNA-encoded OXPHOS signatures, whereas SC1 displayed the reciprocal pattern, with activation of mitochondrial stress adaptation, ISR/amino acid stress, and immature/NKT2-like programs accompanied by loss of terminal NKT1 and TCR-responsive signatures (Fig. 4H). Opa1-KO NKT1 cells adopted an SC1-like transcriptional program, whereas Drp1 co-deletion largely restored a WT-like profile across all seven modules (Supplemental Fig. S9).

Despite increased mitochondrial mass (Fig. 3B, J), Opa1-KO NKT1 cells showed reduced expression of mtDNA-encoded respiratory chain subunits spanning complex I (mt-Nd1, mt-Nd2, mt-Nd4, and mt-Nd5), complex III (mt-Cytb), and complex IV (mt-Co1) (Fig. 4I and Supplemental Fig. S9B). This reduction was accompanied by selective induction of a nuclear encoded mitochondrial maintenance and stress adaption program, including the stress responsive transcription factor Atf5^46^, the biogenesis regulator Ppargc1b^47^, cristae architecture and protein import components (Samm50, Agk, Dnajc11)^48^, matrix chaperones and proteases (Hspa9, Dnaja3, Clpb, Pitrm1, Xpnpep3, Afg3l2)^49^, the ubiquitin ligase Marchf5^50, 51^, the redox regulator Txnrd2, and mitochondrial RNA processing and the mitochondrial translation factors (Rexo2, Elac2, Mrps27, Ptcd3, Gfm1 and Gfm2)^52^ (Fig. 4I and Supplemental Fig. S9B). Thus, Opa1 deficiency leads to an enlarged but transcriptionally compromised mitochondrial compartment accompanied by induction of pathways supporting organelle maintenance, proteostasis, RNA metabolism and mitochondrial translation. At the protein level, ATF5 was increased in Opa1-KO NKT1 cells and partly normalized by Drp1 co-deletion, supporting mitochondrial stress-response program identified by bulk and single-cell transcriptomics (Fig. 4J, K).

In addition, Opa1 deficiency induced a broad integrated stress response and an amino acid metabolic program^42^. Canonical stress response genes Trib3, Asns, Asnsd1, and the translational regulator Eif4ebp1 were increased. This was accompanied by increased expression of genes involved in amino acid transport and sensing (*Slc1a4, Slc3a2, Slc7a1, Slc7a5, Slc7a6,* and *Slc38a9*), serine synthesis and one-carbon metabolism (*Psat1, Shmt2, Mthfd2, Mthfd1l, Mthfs,* and *Chdh*)^53^, and broader amino acid, methionine, and folate metabolism (*Aldh18a1, Bckdhb, Ahcy,* and *Ggh*). A coordinated increase in aminoacyl-tRNA synthetases was also observed (15 of 19 detected cytoplasmic aaRS genes; Fig. 4I and Supplemental Fig. S9B). Together, these changes are consistent with the activation of the integrated stress response^43^.

In parallel, Opa1 deficiency suppressed the transcriptional programs associated with terminal NKT1 differentiation while reinforcing immature/NKT2-like identity (Fig. 4I and Supplemental Fig. S8B)^54^. This was accompanied by coordinated attenuation of TCR responsive immediate early transcription (TCR IEG), including the nuclear receptors (*Nr4a1, Nr4a2,* and *Nr4a3*)^55^, AP-1 family members (*Fos, Fosb, Fosl2, Jun, Junb,* and *Jund*)^56^, and additional immediate early genes (*Dusp1, Dusp5, Dusp10*, *Ier2*, and *Ier5*)^57^. Likewise, an NF-κB transcriptional module comprising NF-κB target genes involved in pathway activation, negative feedback, and survival (*Rel, Relb, Nfkb2, Nfkbia, Nfkbiz, Tnip1, Tnfaip3, Birc2,* and *Birc3*)^58^ was broadly suppressed. Across these mitochondrial, stress response, differentiation, and signaling modules, Drp1 co- deletion shifted expression toward the WT pattern, consistent with restoration of a mature, TCR responsive NKT1 transcriptional state (Fig. 4G).

### Opa1 deficiency shifts iNKT cells toward glycolytic dependence

Bulk RNAseq suggested that Opa1 deficiency remodeled glucose metabolism (Supplemental Fig. S7). *Txnip* encodes a negative regulator of glucose uptake that promotes internalization of GLUT1^59^ and is downregulated in both Opa1-KO NKT1 and NKT2 (Supplemental Fig. S7). Together with induction of *Pfkp* in NKT1 cells and *Hk2* and *Ldhb* in NKT2 cells, these findings suggested remodeling of glucose metabolism in Opa1-KO iNKT cells. We therefore directly assessed metabolic dependency using SCENITH, a flow cytometry-based assay that quantifies the relative contribution of glycolysis and mitochondrial oxidative phosphorylation to protein synthesis^60^. We first validated this approach across thymic T cell populations (Supplemental Fig. S10). Anti-puromycin staining closely paralleled Ki67 expression (Supplemental Fig. S10A, B), indicating that puromycin incorporation accurately reflects the expected hierarchy of translational and proliferative activity among developing thymocytes. SCENITH revealed metabolic reprogramming following Opa1 deletion, characterized by increased glucose dependency and reduced mitochondrial dependency (Fig. 5A, B and Supplemental Fig. S10C-J). The shift was modest in conventional CD4 and CD8 T cells but pronounced across all three iNKT cell subsets (Fig. 5A, B, Supplemental Fig. S10C-J). Consistent with these functional measurements, GLUT1 expression was selectively increased in Opa1-deficient iNKT cell subsets but remained largely unchanged in conventional T cells (Fig. 5C and Supplemental Fig. S9K-N). Concurrent Drp1 deletion shifted both glucose and mitochondrial dependency together with GLUT1 expression toward WT levels across all iNKT subsets, indicating that the metabolic reprogramming results from the fusion-fission imbalance created by Opa1 deficiency.

**Figure 5.**
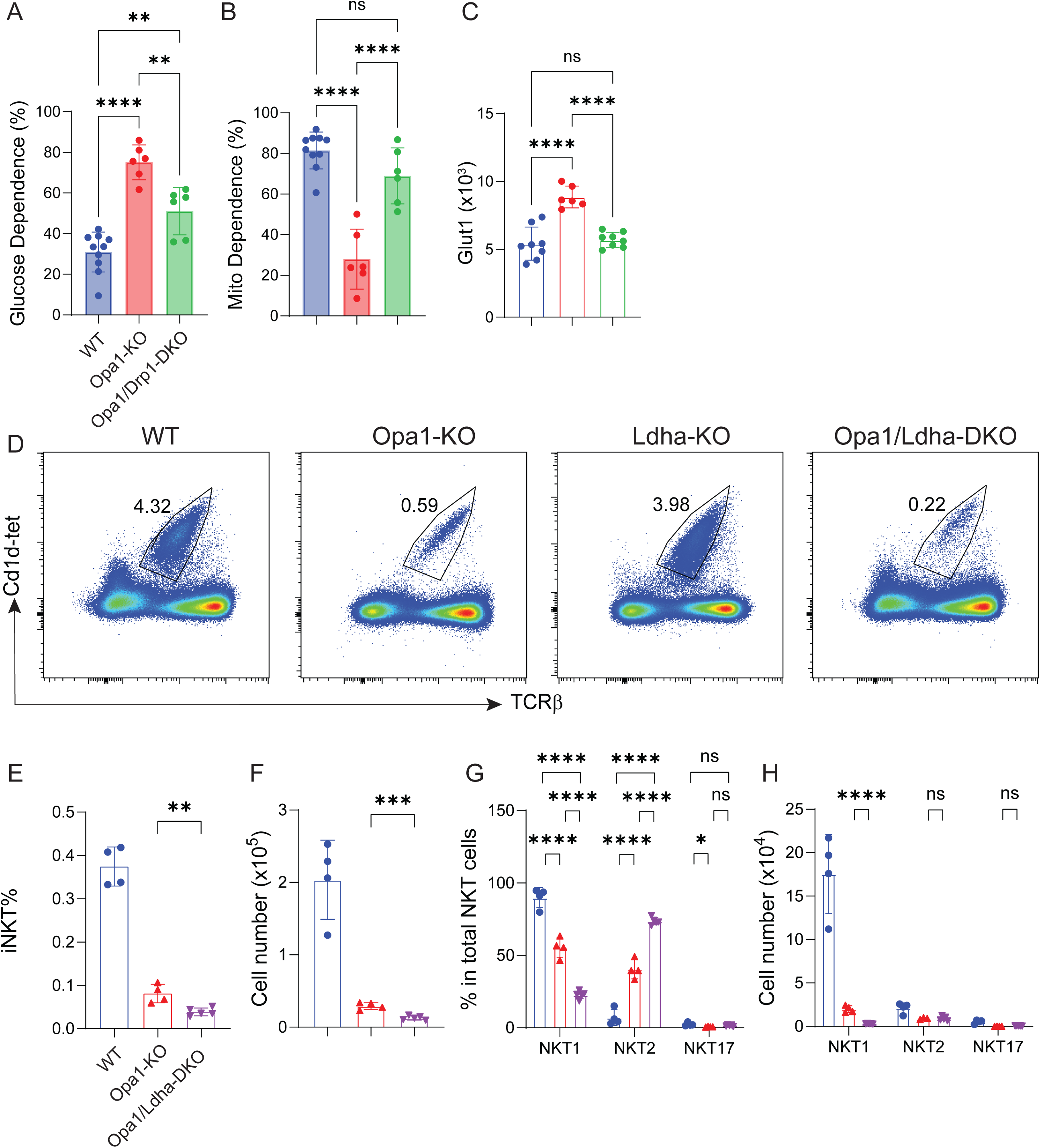
Opa1 deficiency induces compensatory glycolytic remodeling that becomes functionally required for iNKT cell development. (A, B) Glucose dependence (A) and mitochondrial dependence (B) of thymic NKT1 cells from WT, Opa1-KO, and Opa1/Drp1-DKO mice, determined by SCENITH. (C) Surface GLUT1 expression (gMFI) on thymic NKT1 cells of the indicated genotypes. (D) Representative flow cytometry of thymic iNKT cells (CD1d- tetramer⁺TCRβ⁺) from WT, Opa1-KO, Ldha-KO, and Opa1/Ldha-DKO mice; numbers indicate the percentage of cells within CD8α^-^ thymocytes. (E, F) Frequency (E) and absolute number (F) of thymic iNKT cells. (G, H) Percentage within total iNKT cells (G) and absolute number (H) of NKT1, NKT2, and NKT17 subsets. Representative results from three independent experiments were shown. Bars represent mean ± SD. Statistical significance was determined by one-way ANOVA test. ns, not significant; *P < 0.05; **P < 0.01; ***P < 0.001; ****P < 0.0001.

To determine if this metabolic adaptation contributes functionally to iNKT cell development, we genetically ablated the glycolytic enzyme LDHA. Ldha deletion alone had no detectable effect on total thymic iNKT cell frequency, cell number, or subset composition (Supplemental Fig. S11A-D), indicating that LDHA is dispensable for steady-state iNKT-cell development. In contrast, combined deletion of Opa1 and Ldha further reduced total iNKT cell frequency and cellularity compared with Opa1 deficiency alone (Fig. 5D-F). Moreover, Opa1/Ldha DKO mice exhibited a more severe differentiation defect, characterized by further reduction of NKT1 cell percentage and cellularity with reciprocal increase of NKT2 cell percentage among total iNKT cells, whereas NKT17 cells were largely unchanged (Fig. 5G, H). Conventional CD4 and CD8 T cell development remained unaffected in Opa1/Ldha DKO mice (Supplemental Fig. S11E-H), demonstrating that the genetic interaction is selective for the iNKT cell lineage. These findings indicate that the glycolytic remodeling induced by Opa1 deficiency serves as a compensatory metabolic adaptation that partially sustains iNKT cell differentiation.

### Opa1 loss dysregulates mitochondrial DNA content, calcium homeostasis, and TCR signaling

Given the reduction in mtDNA-encoded respiratory chain transcripts in Opa1-KO iNKT cells (Fig. 4I, Supplemental Fig. S7, S9B), we next examined mitochondrial DNA (mtDNA) content. Quantitative PCR analysis of the mtDNA-to-nuclear DNA ratio (*mt-Co1/Tert*) demonstrated that Opa1 deficiency significantly reduced mtDNA content in both conventional CD4 T cells and NKT1 cells, although the defect was substantially greater in iNKT cells (Fig. 6A, B). Concomitant deletion of *Drp1* completely restored mtDNA content in CD4 T cells but only partially restored it in iNKT cells. These findings are consistent with mtDNA depletion contributing to the reduced expression of mtDNA-encoded respiratory chain genes in Opa1-deficient iNKT cells (Fig. 4I, Supplemental Fig. S9B).

**Figure 6.**
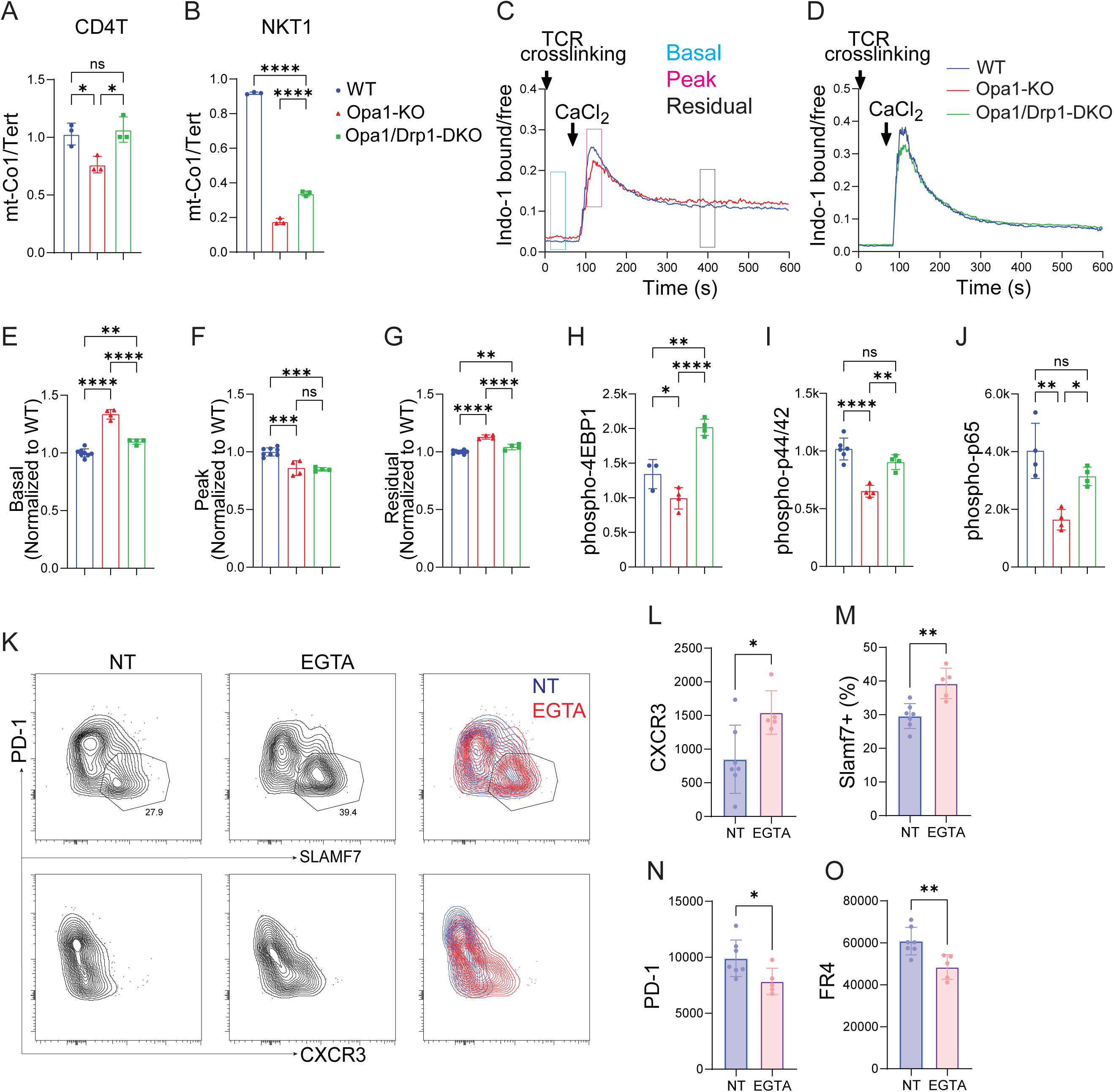
Opa1 loss disrupts mitochondrial DNA content, calcium homeostasis, and TCR proximal signaling in iNKT cells. (A, B) Relative mitochondrial DNA (mtDNA) content in thymic conventional CD4 T cells (A) and NKT1 cells (B), determined by quantitative PCR as the mt-Co1/Tert ratio. (C, D) Representative Indo-1 traces of TCR induced calcium responses in thymic iNKT cells. Cells were stimulated by TCR crosslinking in Ca2+-free medium, followed by CaCl₂ addition. Boxes in (C) indicate the intervals used to quantify basal, peak, and residual calcium. WT and Opa1-KO (C) or WT and Opa1/Drp1-DKO (D) thymocytes were differentially labeled with CFSE and analyzed within the same sample. (E-G) Quantification of basal (E), peak (F), and residual (G) Indo- 1 bound/free ratios, each normalized to paired WT cells analyzed in the same tube. (H-J) Phosphorylation of 4EBP1 (H), ERK1/2 (p44/42) (I), and p65 (J) in thymic NKT1 cells, measured by phospho-flow cytometry 90 min after αGalCer administration. (K) Representative flow cytometry of PD-1 versus SLAMF7 (upper) and PD-1 versus CXCR3 (lower) in iNKT cells from Opa1-KO neonatal thymic organ cultures treated with vehicle (NT) or EGTA; overlays at right. Numbers indicate the percentage of cells within the gate. (L-O) Expression of CXCR3 (L), SLAMF7 (M), PD-1 (N), and FR4 (O) in Opa1-KO iNKT cells after 7 days of culture. Each symbol represents an individual replicate (A, B), an individual time point (E-J) or an individual thymic lobe (L-O); data are mean ± SD, representative from 3 independent experiments. Statistical significance was determined by one-way ANOVA test (A, B, E-J) or unpaired two-tailed Student’s t test (L-O). ns, not significant; *P < 0.05, **P < 0.01, ***P < 0.001, ****P < 0.0001.

Because Opa1 deficiency impaired mitochondrial calcium uptake (Fig. 3L-O), we next asked whether cytosolic calcium homeostasis was also altered. WT and Opa1-KO thymocytes (or WT and Drp1-KO, or WT and Opa1/Drp1-DKO) were differentially labeled with CFSE, mixed within the same sample, enriched for iNKT cells, and loaded with the ratiometric calcium indicator Indo-1. Cells were stimulated by TCR crosslinking in Ca²⁺-free medium followed by extracellular CaCl₂ addition to induce store-operated calcium entry (SOCE)^61^ (Fig. 6C, D). Calcium responses were normalized to paired WT cells analyzed in the same tube (Fig. 6E-G). Drp1 deletion alone did not alter calcium responses (Supplemental Fig. S12A-D). Prior to CaCl₂ addition, Opa1-KO iNKT cells displayed elevated basal cytosolic Ca²⁺ compared with WT cells (Fig. 6E). Following SOCE induction, Opa1-KO cells exhibited a reduced peak calcium response together with persistently elevated cytosolic Ca²⁺ during the plateau phase of the response (Fig. 6F, G). Reciprocal CFSE labeling yielded comparable results (Supplemental Fig. S12E-H). Concomitant deletion of Drp1 partially normalized basal and residual cytosolic Ca²⁺ but failed to restore the SOCE peak (Fig. 6E-G). Together, Opa1 deficiency disrupts cytosolic calcium homeostasis rather than simply attenuating TCR-induced calcium signaling, consistent with impaired mitochondrial calcium handling (Fig. 3L-O).

IPA and module scoring of the transcriptomic data suggested suppression of multiple TCR signaling pathways, including AKT/mTOR, ERK/MAPK, and NF-κB signaling (Fig. 4D,H). To validate these predictions at the protein level, mice were injected intravenously with αGalCer, and phosphorylation of 4EBP1, ERK1/2 (p42/44), and p65 was quantified in thymic NKT1 cells 90 min later by phospho-flow cytometry (Fig. 6H-J). Opa1-KO NKT1 cells exhibited significantly reduced phosphorylation of all three signaling molecules compared with WT controls, whereas concomitant deletion of Drp1 restored each pathway to near WT levels (Fig. 6H-J). Notably, this recovery occurred despite persistent impairment of the SOCE peak (Fig. 6C-G). This dissociation suggests that distinct components of TCR-responsive signaling differ in their sensitivity to OPA1/DRP1-dependent mitochondrial perturbation.

To determine whether dysregulated calcium homeostasis contributes functionally to the developmental defect, we used a neonatal thymic organ culture system^13^, which recapitulated the selective impairment in NKT1 differentiation observed in vivo (Supplemental Fig. S12I-O). Conventional CD4 and CD8 T cell development was preserved, whereas Opa1-KO cultures showed reduced iNKT cell frequency together with decreased expression of the NKT1 markers CXCR3 and SLAMF7 and increased expression of the NKT2 markers PD-1 and FR4 (Supplemental Fig. S12I-O). Reducing extracellular Ca²⁺ with EGTA did not alter the frequency of conventional T cells or total iNKT cells (Supplemental Fig. S12P-R) but increased CXCR3 and SLAMF7 expression while reducing PD-1 and FR4 expression in Opa1-KO iNKT cells (Fig. 6K-O). These data suggest that altered calcium homeostasis contributes functionally to the impaired terminal differentiation of Opa1- deficient iNKT cells. Together with the partial recovery of cytosolic calcium dynamics following Drp1 co-deletion, these findings link mitochondrial remodeling to the calcium homeostasis required for NKT1 differentiation.

## Discussion

Here we identify a genetic interaction between the opposing mitochondrial remodeling proteins OPA1 and DRP1 that controls terminal iNKT cell differentiation. Although OPA1 is broadly required for mitochondrial inner membrane organization, its loss produced a disproportionately severe defect in iNKT cells, characterized by failure of terminal NKT1 differentiation and persistence of an immature/NKT2-like state (Fig. 2)^36^. This phenotype was accompanied by altered mitochondrial morphology, impaired membrane potential and calcium uptake, mtDNA depletion, mitochondrial stress and ISR activation, attenuated TCR-responsive signaling, and a shift toward glycolytic dependence. Importantly, co-deletion of DRP1 substantially improved many of these defects and restored NKT1 subset distribution and phenotypic maturation. These findings argue that iNKT cell differentiation depends not simply on OPA1 as an isolated mitochondrial protein, but on an appropriate balance of mitochondrial membrane remodeling during development.

We previously found that thymic NKT2 cells have higher mitochondrial content and activity than NKT1 cells, consistent with their higher proliferative state^13^. Here, Opa1 deficiency reduced mitochondrial dependency across all three iNKT cell subsets (Fig. 5, Supplemental Fig. S10) and induced mitochondrial stress and ISR associated transcriptional programs in both NKT1 and NKT2 cells (Supplemental Fig. S7). Nevertheless, NKT2 cells were relatively preserved, whereas terminal NKT1 cell differentiation was markedly impaired (Fig. 2). In parallel, deletion of Opa1 with Tbet-cre which targets Tbet expressing cells during NKT1 cell differentiation^37^, did not reproduce the severe NKT1 defect seen with CD4-Cre (Supplemental Fig. S2). One possibility is that the mitochondria-rich precursor/NKT2 state has greater capacity to tolerate impaired mitochondrial quality or excessive fragmentation, whereas progression toward mature NKT1 cells requires a more tightly controlled mitochondrial checkpoint. The recurrent vulnerability of NKT1 cells across metabolic perturbations may reflect the developmental timing of that checkpoint rather than an ongoing requirement in differentiated cells^7–9, 62–65^.

Mitochondrial remodeling may influence this developmental transition in part through its control of Ca²⁺ homeostasis^66^. Shifting mitochondria toward fission by MFN2 depletion reduces mitochondrial Ca²⁺ uptake and retention capacity and impairs store-operated Ca²⁺ entry, whereas inhibiting DRP1 increases mitochondrial Ca²⁺ uptake and accelerates SOCE^67^. Similarly, suppressing MFF in neurons enlarges mitochondria and increases their Ca²⁺ buffering capacity, thereby reducing activity induced cytosolic Ca²⁺ accumulation^68^. These observations indicate that mitochondrial shape itself can influence the partitioning and kinetics of intracellular Ca²⁺. Consistent with this concept, OPA1-dependent mitochondrial remodeling has been repeatedly linked to Ca²⁺ homeostasis, with consequences for cell differentiation and function in cardiomyocytes^69^, endothelial cells^70^, and neurons^71^. In developing thymocytes, early Opa1 deletion increases maximal cytosolic Ca²⁺ responses, while acute OPA1 inhibition in Jurkat cells increases both maximal and residual cytosolic Ca²⁺ following TCR stimulation^20^. In Opa1-deficient iNKT cells, we observed impaired mitochondrial Ca²⁺ uptake together with elevated basal and residual cytosolic Ca²⁺, but also a reduced SOCE peak (Fig. 3, Fig. 6), indicating dysregulated Ca²⁺ homeostasis rather than a uniform increase or decrease in Ca²⁺ signaling. Drp1 co-deletion partially normalized basal and residual cytosolic Ca²⁺ despite incomplete recovery of the SOCE peak, linking these abnormalities to the altered mitochondrial state produced by Opa1 deficiency. Moreover, reducing extracellular Ca²⁺ during thymic organ culture shifted Opa1-deficient cells toward an NKT1 phenotype (Fig. 6), supporting a functional contribution of altered Ca²⁺ homeostasis to the differentiation defect, although precisely which Ca^2+^ parameter is instructive requires further investigation.

OPA1 deficiency also elicited a prominent mitochondrial stress response. Opa1-KO iNKT cells upregulated a program encompassing mitochondrial proteostasis, RNA processing and translation, together with increased ATF5 expression (Fig. 4, Supplemental Fig. S9). A second stress associated program was enriched for amino acid metabolism and tRNA charging, features characteristic of the integrated stress response (ISR) (Fig. 4, Supplemental Fig. S9). Mammalian mitochondrial stress responses are increasingly recognized as heterogeneous and context dependent. Acute mitochondrial proteotoxic stress can induce mitochondrial chaperones together with extensive remodeling of mitochondrial RNA processing and translation^52^, whereas diverse mitochondrial perturbations converge on an ATF4-dependent ISR that coordinates amino acid, one-carbon and redox metabolism^43^. The temporal organization of these responses may also differ with the nature and severity of mitochondrial dysfunction, with ATF5, ATF4-associated metabolic programs, and mitochondrial proteostasis pathways engaged at distinct stages^72^. Although ATF4 protein was not detectably increased in Opa1-KO iNKT cells, the transcriptional overlap with established ATF4-responsive programs^43^ suggests engagement of an ISR-like state (Fig. 4, Supplemental Fig. S9). Importantly, such stress responses can be adaptive rather than simply manifestations of mitochondrial damage. OMA1-DELE1-dependent ISR signaling preserves glutathione metabolism and limits ferroptosis following respiratory-chain dysfunction^73^, and OPA1 deficiency itself can induce an ATF4-associated program that enhances resistance to oxidative lipid damage^74, 75^. Similarly, ATF4 promotes survival and proliferation of Opa1-deficient neural stem cells^76^, indicating that stress-response programs can help cells accommodate mitochondrial dysfunction rather than directly causing the resulting cellular phenotype.

Metabolic remodeling is integral to this adaptation. OPA1-deficient CD8 T cells mount an amino acid deficiency response despite increased uptake, catabolizing amino acids to feed the TCA cycle and regenerate NAD⁺ and glutathione^23^. In OPA1-deficient fibroblasts, impaired oxidative glucose metabolism is accompanied by increased glycolytic carbon flow and glutamine-dependent reductive carboxylation^77^, demonstrating that the compensatory pathways engaged following OPA1 loss vary with cellular context. Our findings identified that Opa1 deficiency reduced mitochondrial dependence and increased glucose dependence, while LDHA, which was largely dispensable for normal iNKT cell development, became necessary in the Opa1-KO cells (Supplemental Fig. S11, Fig. 5). However, this compensatory metabolic rewiring was insufficient to support terminal NKT1 differentiation. This parallels the extensive but ultimately inadequate metabolic adaptation of Opa1-deficient effector CD8 T cells^23^. In contrast, Drp1 co-deletion substantially improved mitochondrial function together with stress-associated transcriptional programs, TCR-responsive signaling and NKT1 differentiation (Figs. 3-6). Thus, the developmental consequences of OPA1 deficiency appear to reflect a mitochondrial remodeling state that cannot be fully overcome by stress adaptation or alternative metabolic pathways but can be substantially ameliorated by altering the opposing mitochondrial remodeling machinery.

## Figure Legends

**Supplemental Figure. S1. OPA1 cell-intrinsically promotes terminal iNKT cell differentiation independently of DP thymocyte development or BCL-XL-sensitive survival pathways. (related to Fig. 1D)**

(A-C) Analysis of thymic iNKT cell maturation in WT and Opa1-KO mice using the CD24/CD44/NK1.1 developmental staging scheme. Representative flow cytometry plots (A) and quantification of stage frequencies (B) and absolute cell numbers (C). (D-F) Analysis of thymic iNKT cell subsets based on T-bet and RORγt expression. Representative flow cytometry plots (D) and quantification of subset frequencies (E) and absolute cell numbers (F). (G-L) Mixed bone marrow chimera analysis. Congenically marked WT (CD45.1) bone marrow was mixed 1:1 with WT or Opa1-KO (CD45.2) bone marrow and transplanted into lethally irradiated recipients. Frequencies of donor-derived conventional CD4 T cells (G), CD8 T cells (H), total iNKT cells (I), and NKT1 (J), NKT2 (K), and NKT17 (L) subsets are shown. (M-P) Analysis of thymic double-positive (DP) thymocytes. Representative flow cytometry plots (M) and quantification of DP frequency (N), DP cell number (O), and CD1d expression on DP thymocytes (P). (Q-T) Effect of BCL-XL transgenic expression on thymic iNKT cell development in Opa1-deficient mice. Frequency (Q) and absolute number (R) of total iNKT cells and frequencies (S) and absolute numbers (T) of NKT1, NKT2, and NKT17 subsets. Each symbol represents one mouse. Bars represent mean ± SD, representative from 3 independent experiments. Statistical significance was determined by one-way ANOVA test (B, C, E, F, Q-T), paired two-tailed Student’s t-test (G-L), or unpaired two-tailed Student’s t test (N-P). ns, not significant; *P < 0.05, **P < 0.01, ***P < 0.001, ****P < 0.0001

**Supplementary Figure S2. OPA1 deletion after NKT1 lineage specification does not impair NKT1 cell development. (related to Fig. 1D)**

(A) Immunoblot analysis of OPA1 in sorted NKT1 and conventional CD4 T cells from Opa1-f/f;Tbx21-Cre− (WT) and Opa1-f/f;Tbx21-Cre+ (KO) mice. β-actin was used as a loading control. (B-D) Representative flow cytometry plots (B) and quantification of thymic iNKT cell frequency (C) and number (D). (E-G) Representative NKT1, NKT2, and NKT17 subset gating (E) and quantification of subset frequency among total iNKT cells (F) and cell number (G). Data are mean ± SD; each symbol represents an individual mouse. Statistical significance was determined by unpaired two-tailed Student’s t-test (C, D), or one-way ANOVA test (F, G). ns, not significant.

**Supplemental Figure S3. Mitochondrial membrane potential, mass and calcium uptake in iNKT cell subsets and conventional T cells (related to Fig. 3).**

(A-E) Mitochondrial membrane potential and mass in thymic NKT2 cells (ICOS^hi^CD4^+^) from WT, Opa1-KO and Opa1/Drp1-DKO mice: representative histograms of TMRM (A) and MTG (C) staining, with quantification of TMRM fluorescence (B), MTG fluorescence (D) and the TMRM/MTG ratio (E). (F-J) As in (A-E) for thymic NKT17 cells (ICOS^hi^CD4^-^), (K-O) for thymic CD4^+^ conventional T cells, and (P-T) for thymic CD8^+^ conventional T cells. (U) Representative mitochondrial calcium uptake trace following TCR stimulation using the reciprocal CFSE-labeling strategy. In contrast to the experiment shown in Fig. 3L, Opa1-KO thymocytes were labeled with CFSE and mixed with unlabeled WT thymocytes before Rhod2-AM loading and analysis. Quantification of peak (W) and residual (X) Rhod2-AM signal, each normalized to the pre-stimulation baseline of the same trace and expressed relative to the WT mean within each experiment. Each symbol represents one mouse (B, D, E, G, I, J, L, N, O, Q, S, T) or individual time points (W, X). Data shows one representative experiment from three independent experiments. Bars represent mean ± SD. Statistical significance was determined by one-way ANOVA with Tukey’s multiple-comparison test (B, D, E, G, I, J, L, N, O, Q, S, T) or unpaired two-tailed Student’s t test (W, X). ns, not significant; *P < 0.05, **P < 0.01, ***P < 0.001, ****P < 0.0001.

**Supplemental Figure S4. Single-cell RNAseq quality control and cell filtering. (related to Fig. 4)**

(A-C) Violin plots showing the distributions of detected genes per cell (A), UMI counts per cell (B), and the percentage of mitochondrial reads (C). Plots are shown by genotype (left) and by integrated cluster (right). Red dashed lines indicate the quality control threshold applied during cell filtering. (D) Summary of cell numbers retained throughout quality control, including raw merged cells, cells remaining after quality control filtering, cells removed as doublets by scDblFinder, and the final cell numbers used for downstream analysis.

**Supplemental Figure S5. Canonical marker expressions supporting thymic iNKT cell cluster annotation. (related to Fig. 4A-C)**

(A) Dot plot showing expression of canonical markers used to annotate the ten integrated thymic iNKT cell clusters (C0–C9), grouped as Stage 0 (C0), NKT progenitor (NKTp; C1-C3), NKT2 (C4), NKT17 (C5) and NKT1 (C6-C9). Dot color indicates mean log-normalized expression, Z-score transformed across clusters separately for each gene; dot size represents the percentage of cells within a cluster expressing that gene. (B) Feature plots showing log-normalized expression of representative developmental and subset-associated markers on the integrated UMAP. Cells are plotted in order of increasing expression to improve visualization of sparse positive populations. Color scales are set independently for each gene.

**Supplemental Figure S6. Reference based validation of thymic iNKT cell cluster annotation and cell cycle status. (related to Fig. 4A-C)**

(A) UMAP of thymic iNKT cells from Krovi et al. (GSE152786), reanalyzed independently. Clusters were renumbered (NC0-NC9) to correspond to the cluster identities defined in this study. (B) Developmental module scores derived from the Krovi reference dataset across integrated clusters and genotypes. (C) Proportion of cells assigned to each integrated cluster within each genotype. Error bars indicate 95% bootstrap confidence intervals. (D) G2/M cell-cycle scores across thymic iNKT developmental subsets and genotypes calculated using Seurat CellCycleScoring. Boxes indicate the median and interquartile range; whiskers extend to 1.5× the interquartile range.

**Supplemental Figure S7. Expression of representative Opa1-KO differentially expressed genes across genotypes. (related to Fig. 4D)**

(A, B) Heatmaps showing representative genes differentially expressed between WT and Opa1-KO NKT1 (A) and NKT2 (B) cells. Expression is displayed across WT, Opa1-KO, and Opa1/Drp1-DKO mice. Genes are grouped by functional categories, and values represent row-wise Z-scored mean log-normalized expression (clipped at ±2).

**Supplementary Figure S8. Identification of interferon-responsive and proliferative NKT1 sub clusters. (related to Fig. 4E-H)** Module scores and representative marker gene expression across NKT1 sub-clusters (SC0-SC5), pooled from all four genotypes. Upper panel, UCell module scores for the interferon-stimulated gene (ISG) and proliferation signatures, Z-scored across all cells and averaged within each sub-cluster; full gene lists are provided in Supplementary Table 1. Lower panel, dot plot of representative canonical ISG and cell-cycle associated marker genes. Dot size indicates the percentage of expressing cells; color indicates mean expression Z-scored across sub-clusters. Color scales are capped at ±2.5 (upper) and ±2 (lower).

**Supplementary Figure S9. Transcriptional programs altered by Opa1 deficiency and differentially restored by Drp1 co-deletion in NKT1 cells. (related to Fig. 4H, I)** (A) Mean module scores across genotypes for the transcriptional programs highlighted in Fig. 4. Values represent mean UCell scores after Z-score transformation across all NKT1 cells; fill color is capped at ±1.0. Module gene sets are listed in Supplementary Table 1. (B) Dot plot showing representative genes grouped by transcriptional program across genotypes in NKT1 cells. Dot size indicates the percentage of expressing cells; color indicates mean log-normalized expression, Z-score transformed across genotypes for each gene. (C) Feature plots showing representative genes projected onto the integrated NKT1 UMAP. Color indicates log-normalized expression, scaled independently for each gene.

**Supplemental Figure S10. Validation of SCENITH and metabolic profiling of thymic T cell subsets. (related to Fig. 5)**

(A, B) Ki67 expression (A) and anti-puromycin incorporation (B) across thymic CD4 T, CD8 T, NKT1, NKT2, and NKT17 cells. (C-J) Glucose and mitochondrial dependency of CD4 T (C, G), CD8 T (D, H), NKT2 (E, I), and NKT17 (F, J) cells from WT, Opa1-KO, and Opa1/Drp1-DKO mice, measured by SCENITH. Dependencies were calculated from anti-puromycin GeoMFI as: glucose dependence = 100 × (Co − 2DG)/(Co − DGO) and mitochondrial dependence = 100 × (Co − O)/(Co − DGO), where Co denotes control, 2DG denotes 2-deoxy-D-glucose treatment, O denotes oligomycin treatment, and DGO denotes combined 2DG + oligomycin treatment. (K-N) Surface GLUT1 expression on CD4 T (K), CD8 T (L), NKT2 (M), and NKT17 (N) cells. Data shows one representative experiment from three independent experiments. Bars represent mean ± SD. Statistical significance was determined by one-way ANOVA with Tukey’s multiple-comparison test. ns, not significant; *P < 0.05, **P < 0.01, ***P < 0.001, ****P < 0.0001.

**Supplemental Figure S11. Ldha deficiency does not impair steady state thymic T cell development. (related to Fig. 5)**

(A-D) Frequency (A) and number (B) of total thymic iNKT cells and the frequency (C) and number (D) of NKT1, NKT2, and NKT17 cell subsets in WT and Ldha-KO mice. (E-H) Frequency (E, G) and number (F, H) of thymic CD4 T (E, F) and CD8 (G, H) cells in WT and Opa1/Ldha-CKO mice. Data shows pooled results (A-D) or one representative experiment (E-H) from three independent experiments. Bars represent mean ± SD. Statistical significance was determined by unpaired two-tailed Student’s t test (A, B, E-H), or one-way ANOVA with Tukey’s multiple-comparison test (C, D). ns, not significant.

**Supplementary Figure S12. Calcium flux specificity controls and neonatal thymic organ culture of Opa1-KO iNKT cells. (related to Fig. 6)**

(A-D) Cytosolic calcium flux in thymocytes following TCR crosslinking, comparing WT and Drp1-KO iNKT cells. (A) Representative Indo-1 bound/free ratio traces; (B–D) Quantification of basal (B), peak (C), and residual (D) Indo-1 ratio, each normalized to the WT mean within the same experiment. (E-H) As in (A-D), comparing WT and Opa1-KO iNKT cells. In this experiment, CFSE labeling was reversed relative to Fig. 6C, such that Opa1- KO rather than WT cells carried CFSE. (I-O) Neonatal thymic organ culture of WT and Opa1-KO thymic lobes. Frequencies of CD4⁺ (I), CD8⁺ (J), and iNKT (K) cells among live cells, and expression of CXCR3 (L, gMFI), SLAMF7 (M, % positive), PD-1 (N, gMFI), and FR4 (O, gMFI) on iNKT cells. (P-R) Opa1-KO neonatal thymic organ cultures maintained without (NT) or with EGTA. Frequencies of CD4 T (P), CD8 T (Q), and iNKT (R) cells among live cells. Symbols represent individual timepoint (B-D, F-H), or individual thymic lobe (I-R). Bars indicate mean ± SD. Data are representative of three independent experiments. Statistical significance was determined by unpaired two-tailed Student’s t-test. ns, not significant; *P < 0.05, **P < 0.01, ****P < 0.0001.

## Materials and methods

### Mice

All mouse experiments were approved by Institutional Animal Care and Use Committee at the Oklahoma Medical Research Foundation. The C57BL/6J (B6) (000664), B6.SJL-Ptprca Pepcb/BoyJ (002014), B6;129S- Gt(ROSA)26Sortm1(CAG-COX8A/Dendra2)Dcc/J (018385), B6;CBA-Tg(Tbx21-cre)1Dlc/J (024507), B6(Cg)-Ldhatm1c(EUCOMM)Wtsi/DatsJ (030112, Ldha-flox), B6.Cg-Tg(LCKprBCL2L1)12Sjk/J (013738, Bcl-XL Tg), B6.Cg-Tg(Cd4-cre)1Cwi/BfluJ (022071, CD4-Cre) mice were purchased from The Jackson Laboratory. Opa1-flox and Dnm1l/Drp1-flox mice were generated by H. Sesaki (JHU)^78, 79^ and obtained from R. Pereira (U. of Iowa) and K. Sun (UTHealth Houston) respectively.

### Antibodies and Reagents

The antibodies conjugated with various fluorophores were from BioLegend anti-mouse CD19 (1D3), B220 (RA3-6B2), CD8a (53-6.7), CD4 (GK1.5/RM4-5), TCRb (H57-597), ICOS (C398.4A), CD62L (MEL-14), CD44 (IM7), CD25 (PC61), NK1.1 (PK136), CD24 (M1/69), CD69 (H1.2F3), CXCR3 (CXCR3-173), SLAMF7 (4G2), SLAMF6 (330-AJ), PD-1 (29f.1A12), FR4 (12A5), and from BD Horizon anti-mouse CD49a (Ha31/8), PLZF (R17-809), Tbet (O4-46), RORgt (Q31-378), Opa1 (18/OPA-1). The biotinylated antibodies for iNKT cell enrichment including TER-119 (TER-119), B220 (RA3-6B2), CD19 (6D5), CD8a (53-6.7), CD11b (M1/70), CD11c (N418), F4/80 (BM8), Ly-6G/Ly-6C (RB6-8C5), and γδTCR (GL3), CD62L (MEL-14), and CD24 (M1/69) antibodies, as well as the anti-ARTC2.2 (s+16a) nanobody were from Biolegend. Human and mouse CD1d/PBS-57 tetramers were obtained from the NIH Tetramer Core Facility.

The T cell culture medium, DMEM, and its supplements penicillin/streptomycin, sodium pyruvate, nonessential amino acids, HEPES, 2-mercaptoethanol, as well as live death blue cell stain kit was from ThermoFisher. FBS was from R&D. The 7-aminoactinomycin D (7-AAD) was obtained from BD Biosciences.

PSC833 was from Santa Cruz Biotechnology. Poly-L-ornithine solution (0.1mg/ml) was from ThermoFisher. Fluoromount-G was obtained from Southern Biotech.

### Flow cytometry and dye staining

Single cell suspensions were stained with live death blue stain prior to surface antibodies. For dye staining, the cells were then pretreated with or without 1uM PSC833 in cell culture medium for 10 min prior to addition of mitochondrial dyes, 10nM (final concentration) MitoTracker green FM or 5 nM TMRM (Invitrogen) and incubation for 15 min at 37c.

### Enrichment and isolation of T cells

For all cell sorting experiments, mice were i.v. injected with 50ug anti-ARTC2.2 (s+16a) nanobody before sacrificing^80^. iNKT cells were negatively enriched using biotinylated anti-mouse Ter119, B220, CD19, CD8a, CD11b, CD11c, F4/80, Ly-6G/Ly-6C, γδTCR, CD62L and MojoSort Streptavidin Nanobeads (BioLegend). iNKT cells were sorted as 7-AAD^-^TCRβ^int^CD1dtet^+^ cells; NKT1 cells were sorted as ICOS^lo^ iNKT cells; NKT2 cells were sorted as ICOS^hi^CD4^+^ iNKT cells; naïve CD4 T cells were sorted as 7-AAD^-^CD1dtet^-^TCRβ^+^ CD4^+^CD25^-^ CD62L^+^.

### Proteomics

Freshly sorted naïve CD4 T and iNKT were processed as described previously^81^. Briefly, the cells were washed in PBS once and lysed with protein lysis buffer (150 mM sodium chloride, 1% NP-40, 0.5% sodium deoxycholate, 0.1% sodium dodecyl sulfate, 50 mM Tris, pH7.4) with 1 × Halt Protease and Phosphatase Inhibitor cocktail (Thermo Scientific). Data-independent acquisition proteomics was performed by the IDeA National Resource for Quantitative Proteomics using their standard pipeline (Spectronaut directDIA search against mouse UniProtKB, 1% precursor and protein q-value, maxLFQ inference). Intensities were normalized at the sample level (ProteiNorm). Differential abundance was assessed by limma with empirical Bayes smoothing, with an FDR-adjusted p < 0.05 considered significant. For display, abundance values were row- wise standardized (Z-score) and capped at ±1.2. Heatmaps were generated in R using pheatmap (v1.0.12). The proteomics data has been deposited at Zenodo (https://doi.org/10.5281/zenodo.15233269).

### Mitochondrial DNA quantification by real-time PCR

The thymic CD4 T and iNKT cells were freshly sorted. The genomic DNA and mitochondrial DNA were extracted with DNeasy Blood & Tissue (QIAGEN) according to the manufacturer’s protocol. The real-time PCR was performed using Universal SYBR Green fast qPCR master mix (ABclonal) with respective primers in a LightCycler 480 Instrument II (Roche). The relative mitochondrial DNA/nuclear DNA (mtDNA:nDNA) ratio was calculated using the ΔΔCt method.

mt-CO1_Fwd: GCCCCAGATATAGCATTCCC;

mt-CO1_Rev: GTTCATCCTGTTCCTGCTCC;

Tert_Fwd: CTAGCTCATGTGTCAAGACCCTCTT;

Tert_Rev: GCCAGCACGTTTCTCTCGTT.

### Confocal microscopy analysis

The sorted CD4 T, as well as iNKT cells from Dendra2 transgenic mice were seeded in 96-well glass bottom plates (Cellvis; 0.170±0.005mm) coated with poly-L-ornithine (0.1mg/ml) and the cells were fixed with fixative buffer (2% paraformaldehyde with 0.075% glutaraldehyde in PBS) for 10 min prior to mounting with Fluoromount-G. The Z-stacked images of the mitochondrial were acquired using a Zeiss LSM 880 Confocal Laser Scanning microscope with airyscan using a 100X oil immersion objective with 5X zoom. The images were processed using the Airyscan Processing features in the Zen Black software, then transferred to Imaris software (version 9.3.1) for 3D rendering and analysis. An iso-surface was created around Dendra2 green fluorescence signal with the seed points diameter of 0.3 um (size of an individual mitochondrion based on EM data) was used.

### Calcium flux assay

As described previously^82^,, thymocytes from WT mice were pre-enriched for iNKT cells (see above), labeled with CFSE (10 nM in PBS) for 10 min on ice, washed, and mixed at a 1:1 ratio with unlabeled thymocytes from mutant mice. Reciprocal labeling experiments, in which mutant thymocytes were CFSE-labeled and WT thymocytes were unlabeled, were performed as controls. Cells were then loaded with either 2 μM Indo-1 AM (Thermo Fisher) or 0.2 μM Rhod-2 AM (Thermo Fisher) in RPMI containing 1% FBS and 4 μM probenecid (Thermo Fisher). After washing, cells were stained with CD1d-αGalCer tetramers and antibodies against TCRβ, CD4, and ICOS. For TCR-induced Ca²⁺ flux measurements, cells were additionally incubated on ice with purified hamster anti-mouse CD3ε antibody (10 μg/mL; clone 145-2C11, BioLegend). Cells were resuspended in PBS, and TCR crosslinking was initiated by addition of goat anti-Armenian hamster IgG (H+L) (Jackson ImmunoResearch). Extracellular Ca²⁺ influx was triggered by the addition of CaCl₂ (final concentration 1 mM). Indo-1 responses were quantified as the ratio of violet to blue fluorescence using FlowJo software (Tree Star), whereas Rhod-2 fluorescence was normalized to the baseline fluorescence before stimulation.

### Thymic organ culture

Timed matings were established, and thymic lobes were harvested from neonatal mice (day 0). Thymic lobes were cultured on 0.4-μm pore-size cell culture inserts (Sigma, CLS3450) placed in six-well plates containing 1.6 mL complete DMEM supplemented with 10% FBS. EGTA was added to the culture medium at a final concentration of 0.2 mM beginning 24 h after culture initiation and maintained throughout the remainder of the culture period. Culture medium was replaced daily. At the indicated endpoint, thymic lobes were dissociated into single-cell suspensions and stained with fluorochrome-conjugated antibodies for flow cytometric analysis.

### Electron microscopy

Freshly sorted iNKT cell subsets were immediately fixed. Sample processing was carried out by the UCSD Electron Microscopy Core Facility. Mitochondrial morphology was visualized by transmission electron microscopy. Mitochondrial size and cristae length were quantified using Fiji ImageJ software. For each condition/cell type, a minimum of 10 images, 20 cells, and 100 mitochondria were analyzed.

### Mixed bone marrow chimeras

Total bone marrow cells were prepared from the femurs and tibias of wild-type (CD45.1^+^) or mutant (CD45.2^+^) donor mice, and samples were depleted of mature T cells with anti-Thy1.2 (30-H12; BD). Recipient F1 mice (CD45.1^+^CD45.2^+^) were pretreated with sulfatrim in drinking water 3 days prior, lethally irradiated (900 rads) and received 10 × 10^6^ T cell-depleted bone marrow cells. Chimeras were maintained on drinking water with sulfatrim and analyzed 6 weeks after transplantation.

### SCENITH assay

Cellular metabolic dependency was determined using SCENITH (single-cell energetic metabolism by profiling translation inhibition) as previously described^60^. Freshly isolated thymocytes were incubated for 10 min at 37°C in complete medium containing vehicle, 100 mM 2-deoxy-D-glucose (2-DG), 1 μM oligomycin, or both inhibitors. Puromycin was then added to a final concentration of 10 μg/mL, and cells were incubated for an additional 20 min then stained with fluorochrome-conjugated antibodies against surface markers and a viability dye. Cells were washed, fixed and permeabilized using Fix/Perm buffer (Tonbo), stained with anti-puromycin- PE antibody (clone 2A4, BioLegend), and analyzed by flow cytometry. Glucose dependency and mitochondrial dependency were calculated from the geometric mean fluorescence intensity (gMFI) of puromycin staining as previously described by Argüello et al^60^.

### Bulk RNA sequencing and data analysis

Total RNA was isolated from thymic iNKT cell subsets sorted from littermate mice of different genotypes as noted, using the miRNeasy Micro Kit (QIAGEN). RNA integrity was reflected by RIN>9 across all samples. RNAseq libraries were prepared from 10ng total RNA using the QIAseq UPXome RNA library kit (QIAGEN) according to the manufacturer’s instructions and sequenced with 150bp paired end configuration on NovaSeq X Plus (Illumina).

Sequencing data were analyzed using the QIAseq UPXome analysis workflow in CLC Genomics Workbench (QIAGEN) with the manufacturer’s recommended settings. Briefly, the first 16 bases of Read 2 were removed, read through adapters were trimmed, reads were quality and ambiguity trimmed, 3′ polyA/polyG/polyT homopolymers were removed, and short reads were discarded. Filtered reads were aligned to the *Mus musculus* GRCm39 reference genome in reverse stranded mode, and gene expression was quantified as total read counts, with transcripts per million (TPM) calculated for visualization. Differential expression was assessed with the Differential Expression for RNA-Seq module in CLC Genomics Workbench, which fits a negative binomial generalized linear model to gene level counts and tests differences between groups by Wald test, using TMM normalization with filtering on average expression prior to FDR correction (n = 3-4 samples per group); genes with FDR<0.1 were considered differentially expressed. Differentially expressed genes were analyzed in Ingenuity Pathway Analysis (QIAGEN) for canonical pathway enrichment, with significance assessed by right-tailed Fisher’s exact test (P<0.05). Heatmaps display row wise z-score-transformed log₂(TPM + 1) values, with gene and sample order manually specified.

### Single cell RNA sequencing and analysis

#### Single-cell RNA sequencing

Total thymic iNKT cells were FACS-sorted from WT, Drp1-KO, Opa1-KO, and Opa1/Drp1-DKO mice (5-15 mice pooled per genotype; 6-8 week-old; sex matched) and fixed with the Evercode Cell Fixation v2 kit (Parse Biosciences). Barcoded libraries were prepared with the Evercode WT v2 kit (Parse Biosciences) and sequenced with 150 bp paired-end reads on a NovaSeq X Plus (Illumina).

#### Preprocessing and quality control

Reads were processed with the Parse Biosciences split-pipe pipeline v1.6.3 and aligned with STAR v2.7.11b to the *Mus musculus* GRCm39 reference (Ensembl annotation release 109) to generate per-cell UMI count matrices. Matrices were analyzed in R v4.5.1 with Seurat v5.3.0 (SeuratObject v5.2.0). Mitochondrial content was calculated as the percentage of UMIs mapping to genes with the mt- prefix. Cells were retained if they expressed 500-6,000 genes, had fewer than 60,000 UMIs, and had <10% mitochondrial transcripts. Doublets were identified with scDblFinder v1.24.10 and removed. Cell numbers at each stage and per-genotype doublet rates are reported in Supplementary Fig. S3.

#### Normalization, integration, and clustering

Gene expression counts were log-normalized, and the 2,000 most variable genes were identified using the variance-stabilizing transformation (“vst”) method. Scaled data were subjected to principal component analysis (30 PCs), followed by Harmony (v1.2.4) integration using genotype as the grouping variable. A shared nearest-neighbor graph was constructed using Harmony dimensions 1–20 (*k* = 20), and cells were clustered using the Louvain algorithm at a resolution of 0.5, yielding ten clusters (C0–C9). UMAP visualization was generated using the same 20 Harmony dimensions. Cluster marker genes were identified using Seurat FindAllMarkers (Wilcoxon rank-sum test; positive markers only; genes detected in ≥25% of cells in either group; minimum log₂ fold change of 0.25). NKT1 cells (clusters C6- C9) were subsequently extracted and re-clustered using the same workflow, except that principal component analysis was performed using 20 principal components, yielding six NKT1 sub-clusters (SC0-SC5).

#### Cluster annotation

Cluster identities were assigned based on canonical iNKT developmental markers and independently validated by scoring published thymic iNKT developmental gene modules from Krovi *et al.* (GSE152786, Supplemental Data I) using Seurat AddModuleScore. Mean module scores were calculated for each cluster × genotype group and visualized after row-wise Z-score transformation.

#### Compositional analysis

Cell composition was quantified as the proportion of cells assigned to each category within the relevant parent population. For global analyses, cluster (C0-C9) and developmental subset (stage 0, NKTp, NKT2, NKT17, and NKT1) proportions were calculated within each genotype. For NKT1 re-clustering, genotype composition was calculated within each NKT1 subcluster. Ninety-five percent confidence intervals were estimated by bootstrap resampling of cells within the corresponding parent population (1,000 iterations; seed = 42), and the 2.5th and 97.5th percentiles of the bootstrap distribution were reported.

#### Module scoring

Nine gene modules were scored. Seven modules (mtDNA-encoded OXPHOS, mitochondrial stress/UPRmt, integrated stress response/amino acid stress, NKT1 terminal maturation, NKT2-like retention, TCR immediate-early (Nr4a/AP-1), and NF-κB) were curated from bulk RNA-seq differential expression analyses together with published gene signatures. Two additional modules were included to define NKT1 sub- cluster identity: an interferon-stimulated gene (ISG) module compiled from published signatures and a proliferation module derived from genes specifically enriched in WT sub-cluster SC5. All module gene lists are provided in Supplementary Table 1. Per-cell module scores were calculated using UCell (v2.14.0; *maxRank* = 1500). For visualization, UCell scores for each module were standardized (Z-score) across all NKT1 cells and then averaged within the indicated groups. Heatmaps summarize module scores by NKT1 sub-cluster (all genotypes pooled) and by genotypes. Cell-cycle phase scores were calculated using Seurat CellCycleScoring with the built-in cc.genes.updated.2019 S-phase and G2/M gene sets after conversion to mouse gene symbols. Cell-cycle effects were not regressed during preprocessing. G2/M scores and the proportion of cycling cells were visualized across developmental subsets and genotypes.

#### Dot plots and feature plots

Dot plots display the mean log-normalized expression of each gene within each group after Z-score transformation across groups; dot size represents the percentage of cells expressing the gene. Feature plots display log-normalized gene expression on the UMAP embedding using a common color scale across all panels, with expressing cells plotted above non-expressing cells.

## Acknowledgments and funding sources

We thank the National Institutes of Health Tetramer Core Facility (contract number 75N93020D00005) for providing CD1d tetramers. This work was supported by National Institute of Health Grants GM-147713 and GM-139763 to MZ, GM-137921 to TL, GM-144103 to HS, NIH Instrument Grant S10OD028479 (Cytek Aurora analyzer), and a Presbyterian Health Foundation (PHF) biomedical research grant to MZ and TL.

## Data Sharing Plan

All data supporting the findings of this study will be made publicly available upon publication. Raw and processed bulk RNA-seq and single-cell RNA-seq data will be deposited in the NCBI Gene Expression Omnibus (GEO). Source data underlying the figures and supplementary figures will be provided with the article or deposited in a public repository. Code used for computational and statistical analyses, together with relevant documentation and gene/module lists required to reproduce the analyses, will be made publicly available through GitHub. Additional data and materials will be available from the corresponding author upon reasonable request.

## Author contributions

M.Z., S.S., J.C., R.L., and T.L.L. designed experiments. S.S., J.C., and M.Z. performed experiments. M.Z., S.S., J.C., R.L. T.L.L. and V.C. performed data analysis. R.L., T.L.L., H.S., R.O.P., and K.S. provided key tools and expertise. The manuscript was written by M.Z. The project was supervised by M.Z.

