## Supplemental Figs for "OPA1-dependent mitochondrial remodeling coordinates TCR signaling and metabolic adaptation during iNKT cell differentiation"

Supplemental S1

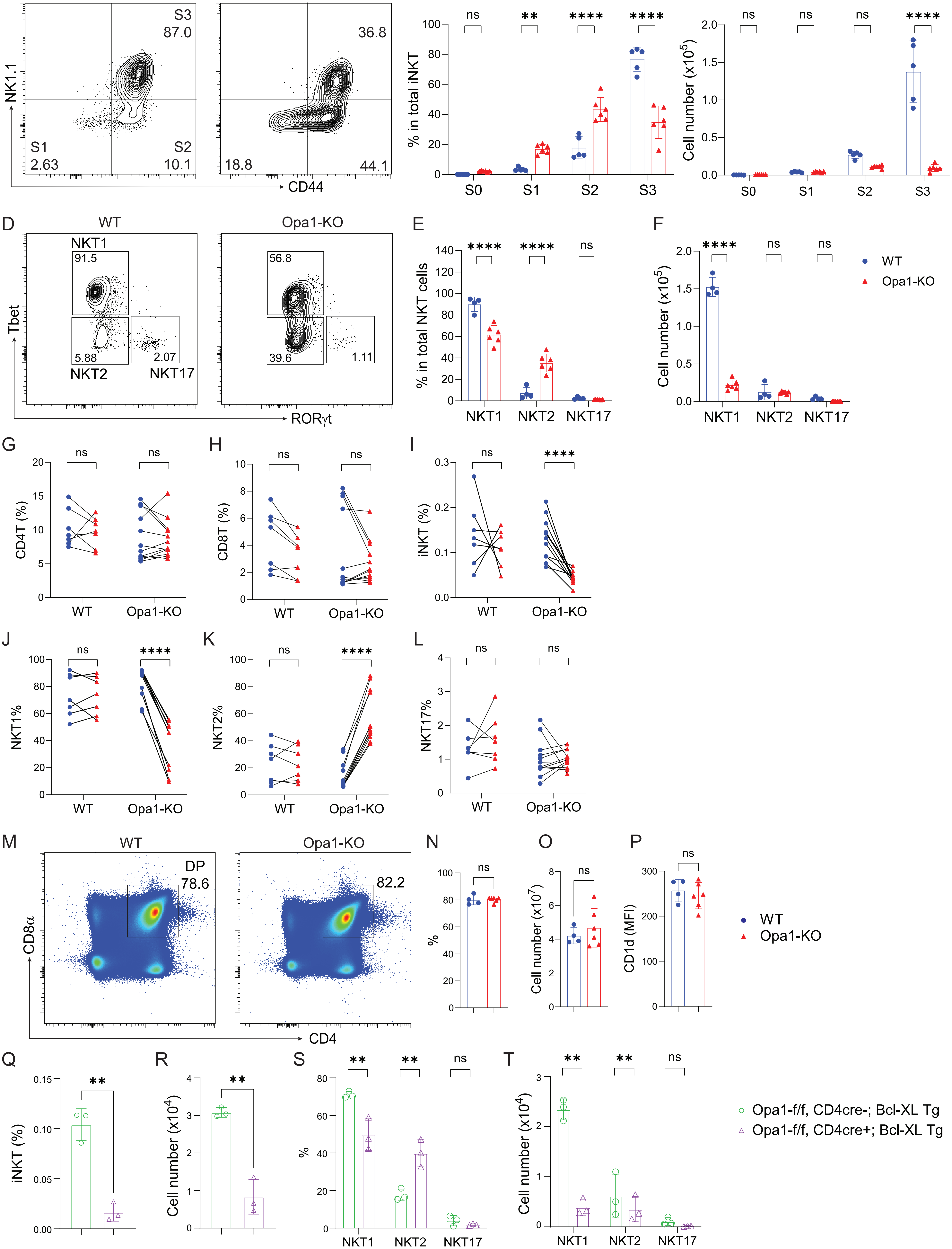

Supplemental S2

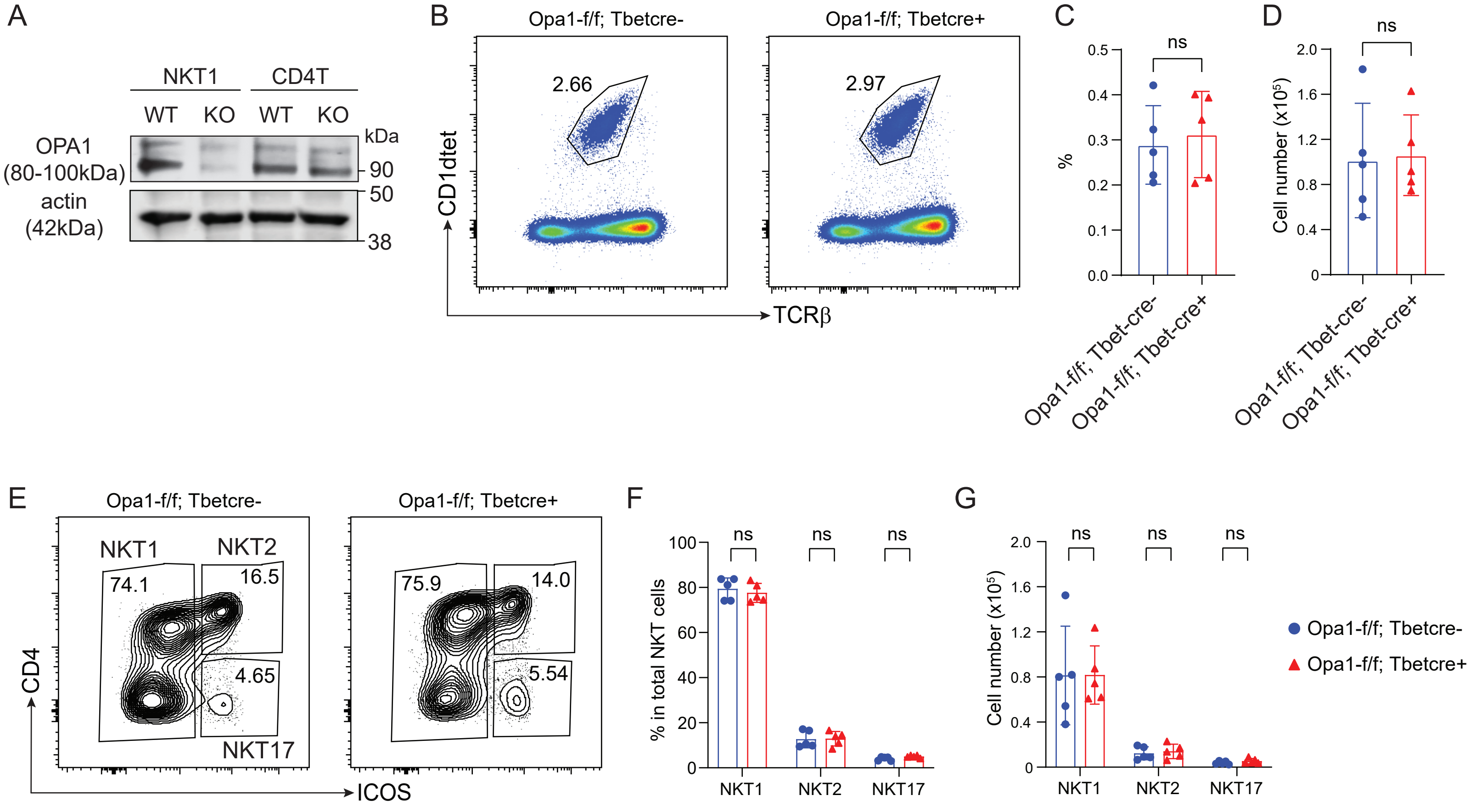

Supplemental S3

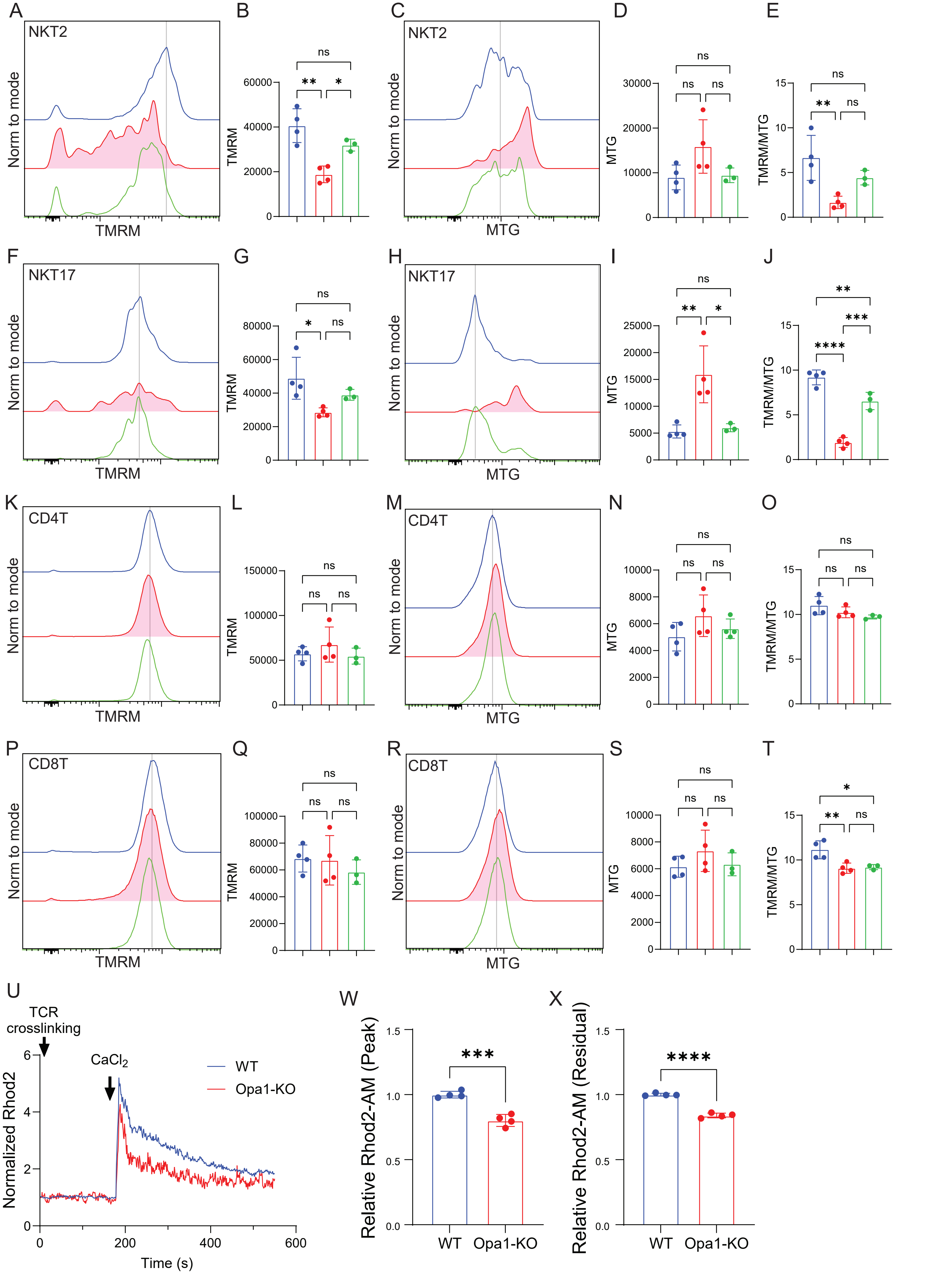

Supplemental S4

A

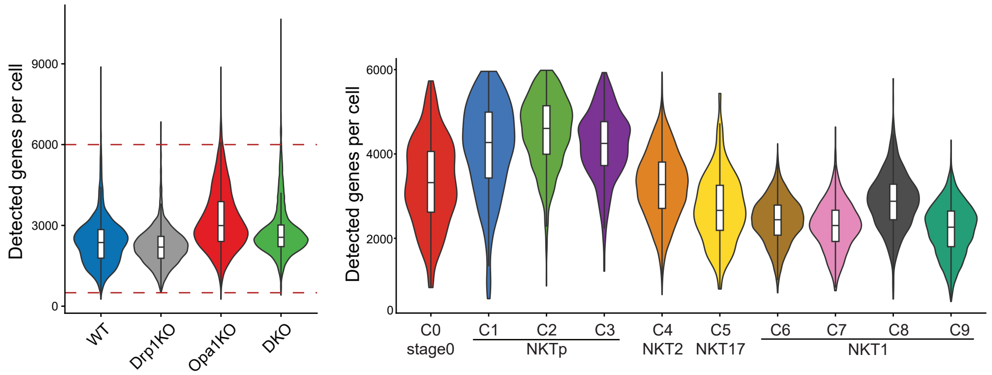


B

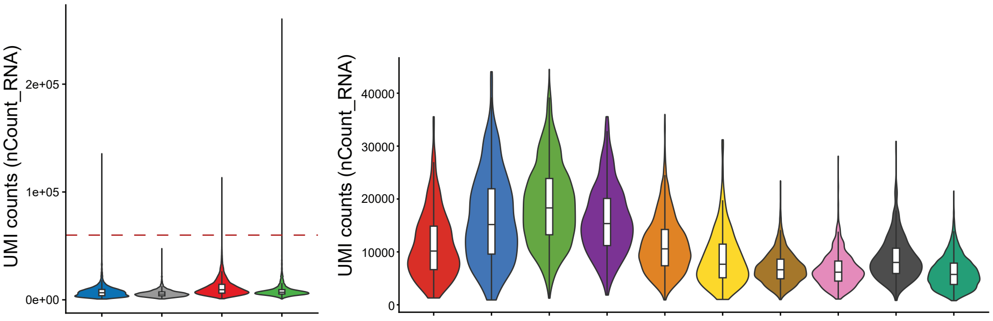


C

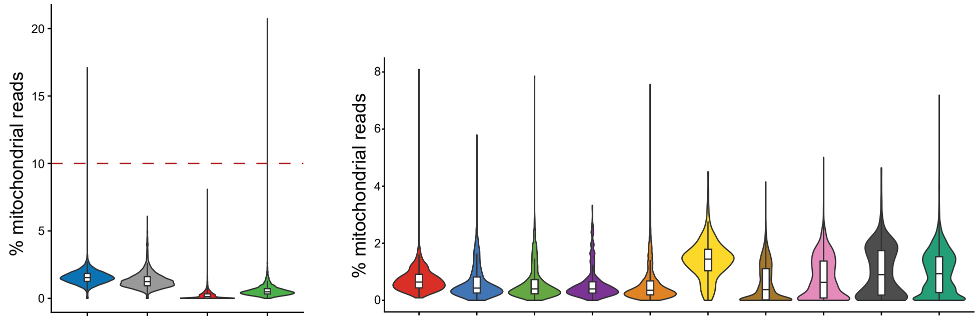


D

| Genotype | Raw (merged) | Post QC | Doublets removed | Post doublet removal | % Doublet | %_retained |
|---|---|---|---|---|---|---|
| WT | 5502 | 5475 | 443 | 5032 | 8.1 | 91.90% |
| Drp1KO | 3292 | 3287 | 194 | 3093 | 5.9 | 94.10% |
| Opa1KO | 9842 | 9690 | 757 | 8933 | 7.8 | 92.20% |
| DKO | 4873 | 4827 | 373 | 4454 | 7.7 | 92.30% |

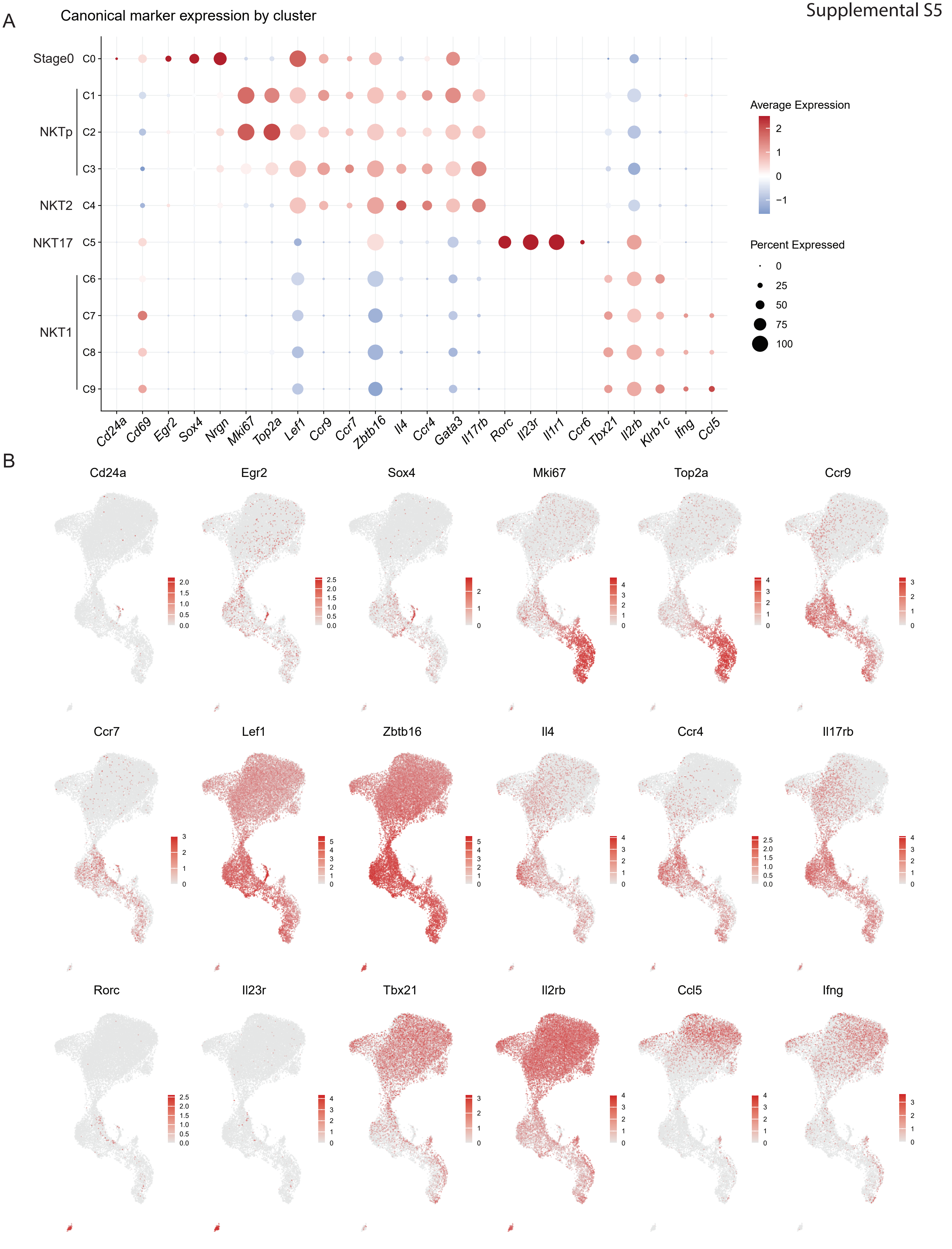
A
Canonical marker expression by cluster
Stage0
NKTp
NKT2
NKT17
NKT1
C0
C1
C2
C3
C4
C5
C6
C7
C8
C9
Cd24a
Cd69
Egr2
Sox4
Nrgn
Mki67
Top2a
Lef1
Ccr9
Ccr7
Zbtb16
Il4
Ccr4
Gata3
Il17rb
Rorc
Il23r
Il1r1
Ccr6
Tbx21
Il2rb
Klrb1c
Ifng
Ccl5
Average Expression
2
1
0
−1
Percent Expressed
0
25
50
75
100
B
Cd24a
Egr2
Sox4
Mki67
Top2a
Ccr9
Ccr7
Lef1
Zbtb16
Il4
Ccr4
Il17rb
Rorc
Il23r
Tbx21
Il2rb
Ccl5
Ifng

Supplemental S6

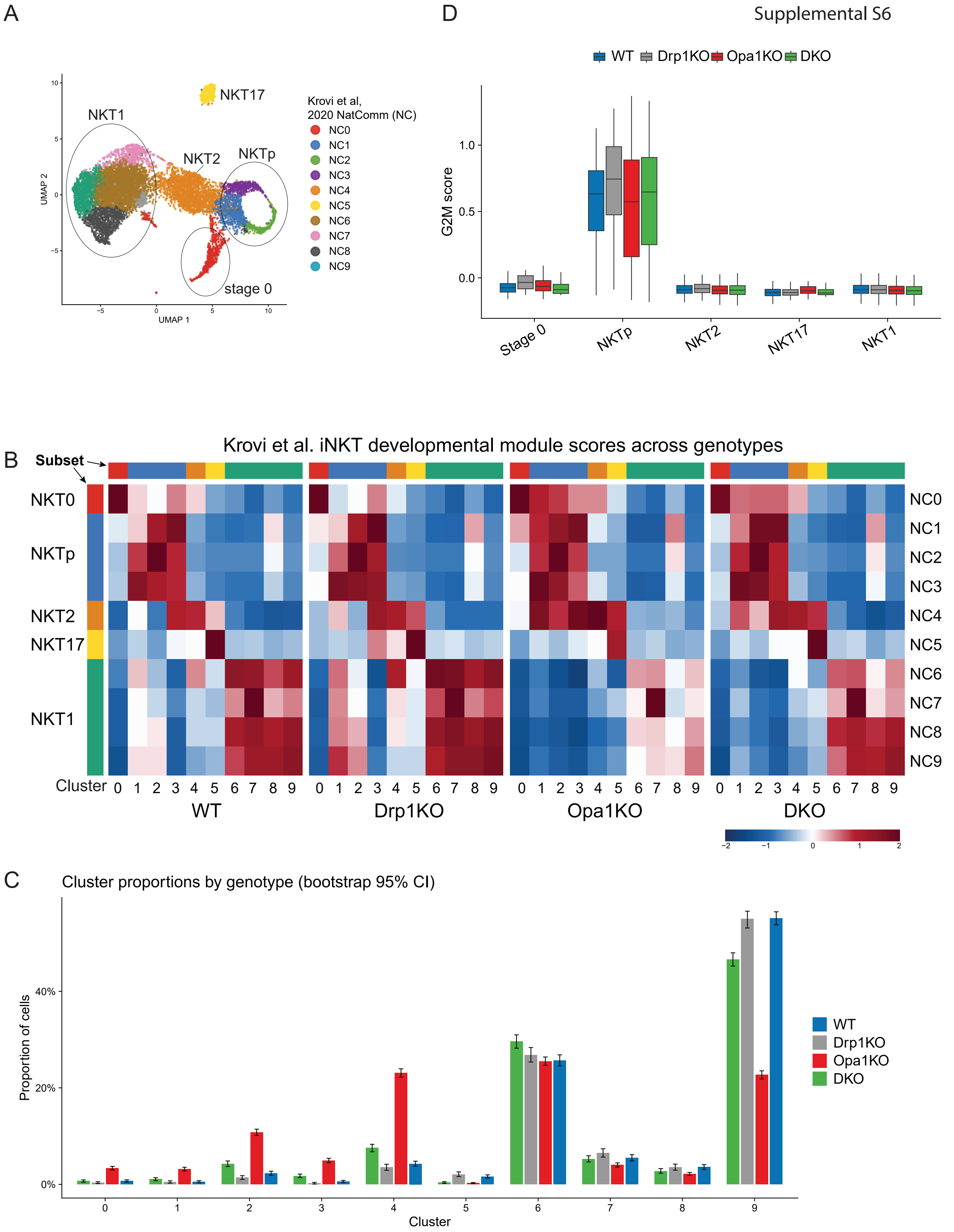

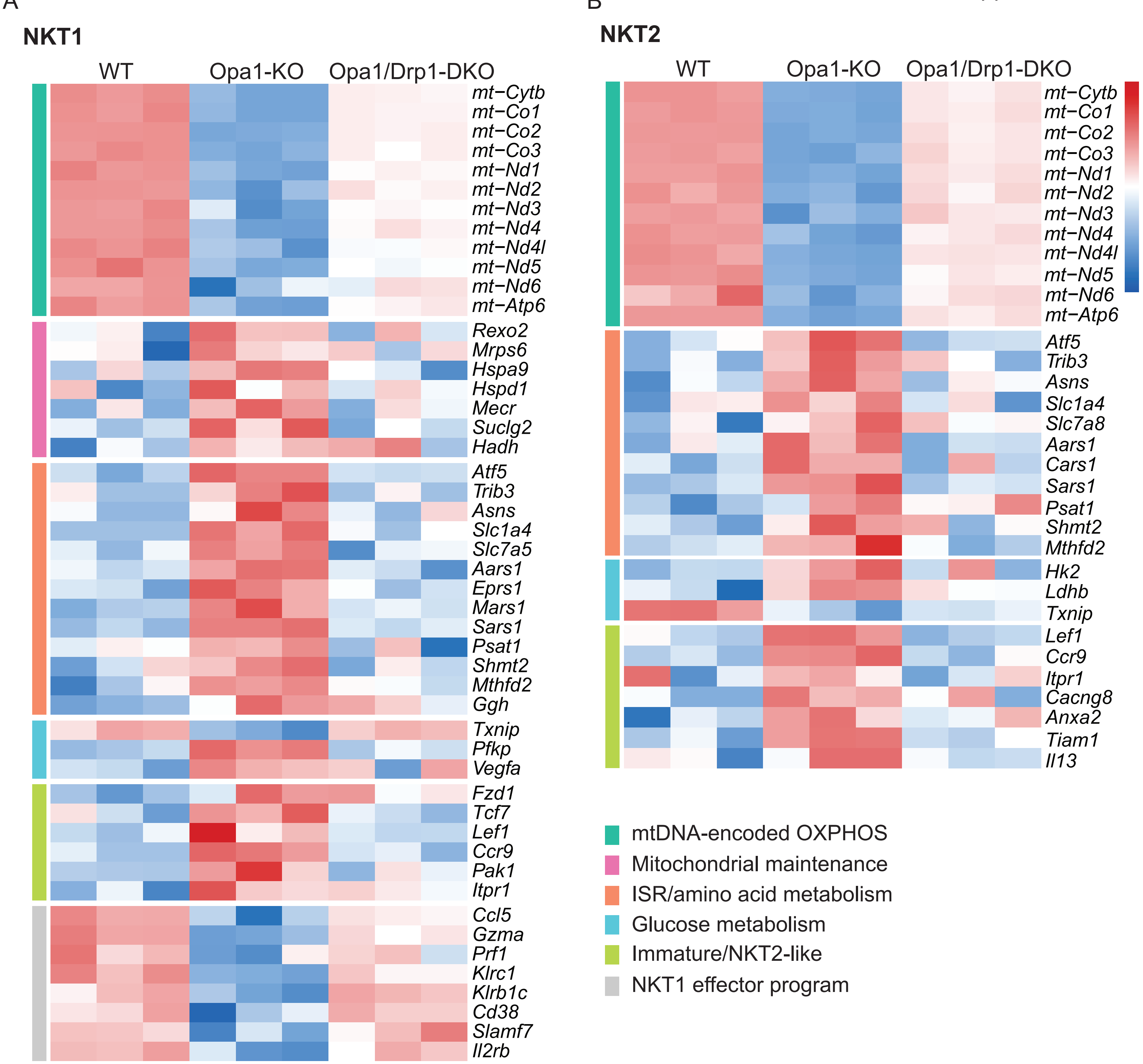
A
NKT1
WT
Opa1-KO
Opa1/Drp1-DKO
mt−Cytb
mt−Co1
mt−Co2
mt−Co3
mt−Nd1
mt−Nd2
mt−Nd3
mt−Nd4
mt−Nd4l
mt−Nd5
mt−Nd6
mt−Atp6
Rexo2
Mrps6
Hspa9
Hspd1
Mecr
Suclg2
Hadh
Atf5
Trib3
Asns
Slc1a4
Slc7a5
Aars1
Eprs1
Mars1
Sars1
Psat1
Shmt2
Mthfd2
Ggh
Txnip
Pfkp
Vegfa
Fzd1
Tcf7
Lef1
Ccr9
Pak1
Itpr1
Ccl5
Gzma
Prf1
Klrc1
Klrb1c
Cd38
Slamf7
Il2rb
B
NKT2
WT
Opa1-KO
Opa1/Drp1-DKO
2
1
0
−1
−2
mt−Cytb
mt−Co1
mt−Co2
mt−Co3
mt−Nd1
mt−Nd2
mt−Nd3
mt−Nd4
mt−Nd4l
mt−Nd5
mt−Nd6
mt−Atp6
Atf5
Trib3
Asns
Slc1a4
Slc7a8
Aars1
Cars1
Sars1
Psat1
Shmt2
Mthfd2
Hk2
Ldhb
Txnip
Lef1
Ccr9
Itpr1
Cacng8
Anxa2
Tiam1
Il13
mtDNA-encoded OXPHOS
Mitochondrial maintenance
ISR/amino acid metabolism
Glucose metabolism
Immature/NKT2-like
NKT1 effector program

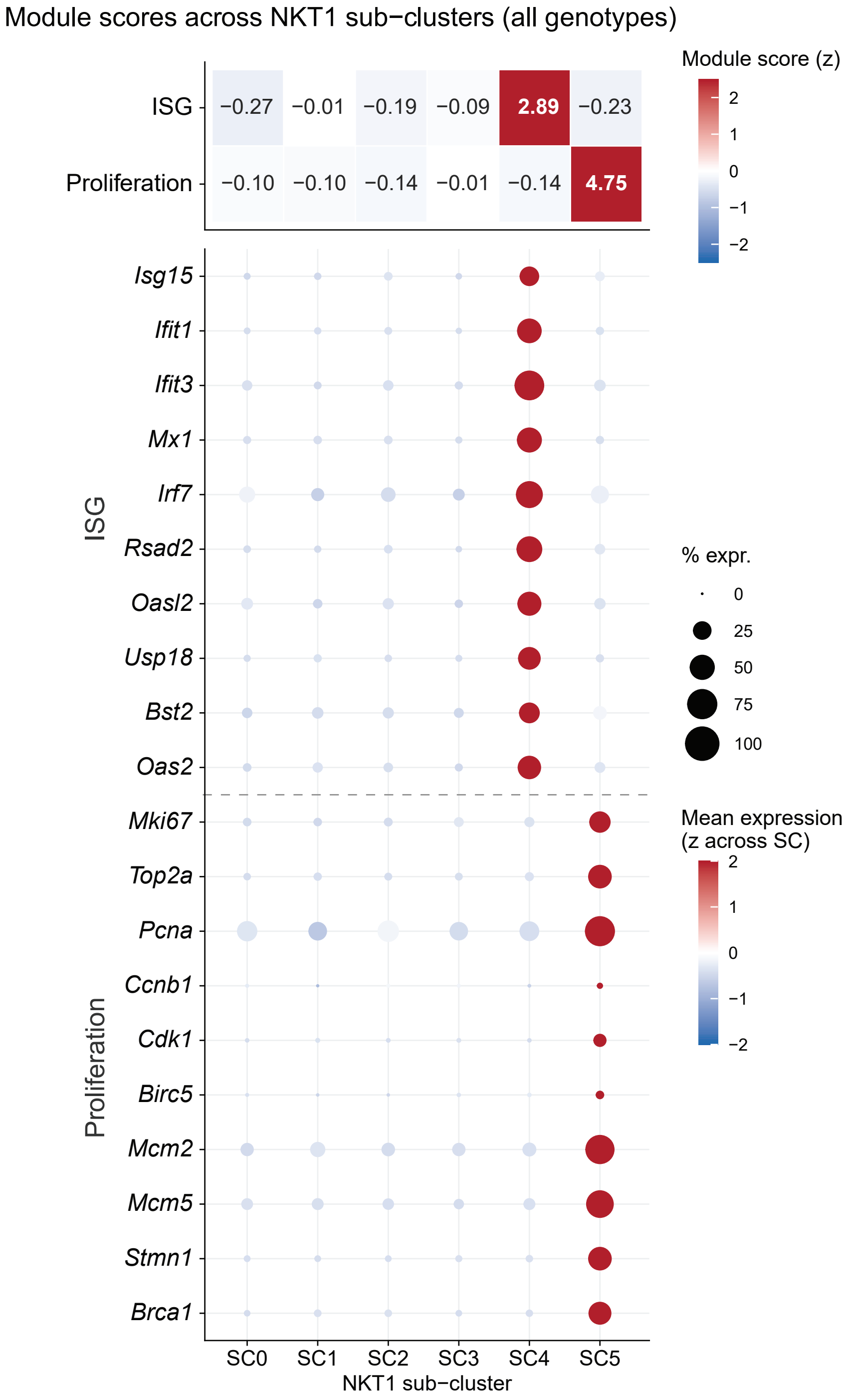
Module scores across NKT1 sub−clusters (all genotypes)
Module score (z)
ISG
Proliferation
−0.27
−0.01
−0.19
−0.09
2.89
−0.23
−0.10
−0.10
−0.14
−0.01
−0.14
4.75
Isg15
Ifit1
Ifit3
Mx1
Irf7
Rsad2
Oasl2
Usp18
Bst2
Oas2
Mki67
Top2a
Pcna
Ccnb1
Cdk1
Birc5
Mcm2
Mcm5
Stmn1
Brca1
SC0
SC1
SC2
SC3
SC4
SC5
NKT1 sub−cluster
% expr.
0
25
50
75
100
Mean expression
(z across SC)

Supplemental S9

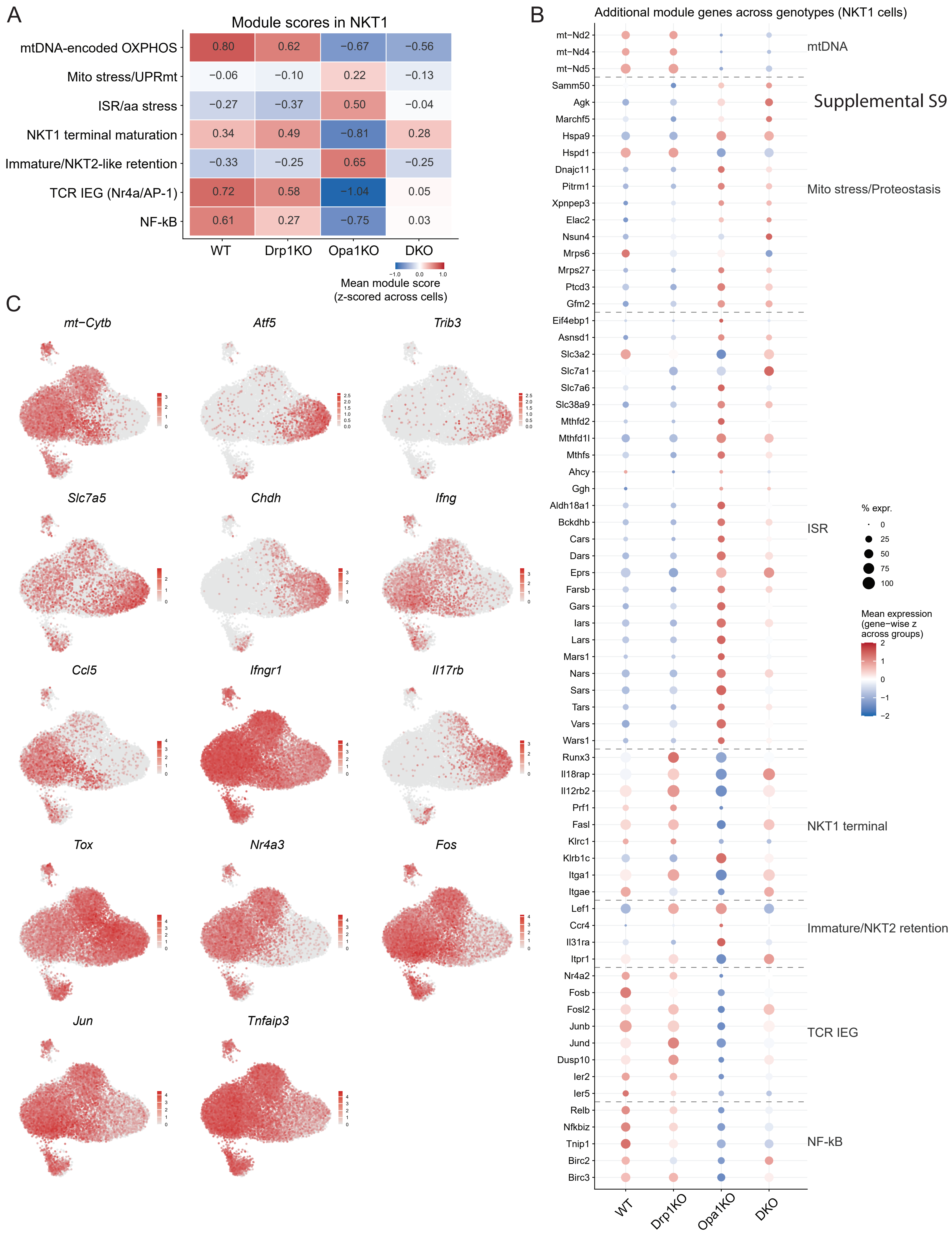

Supplemental S10

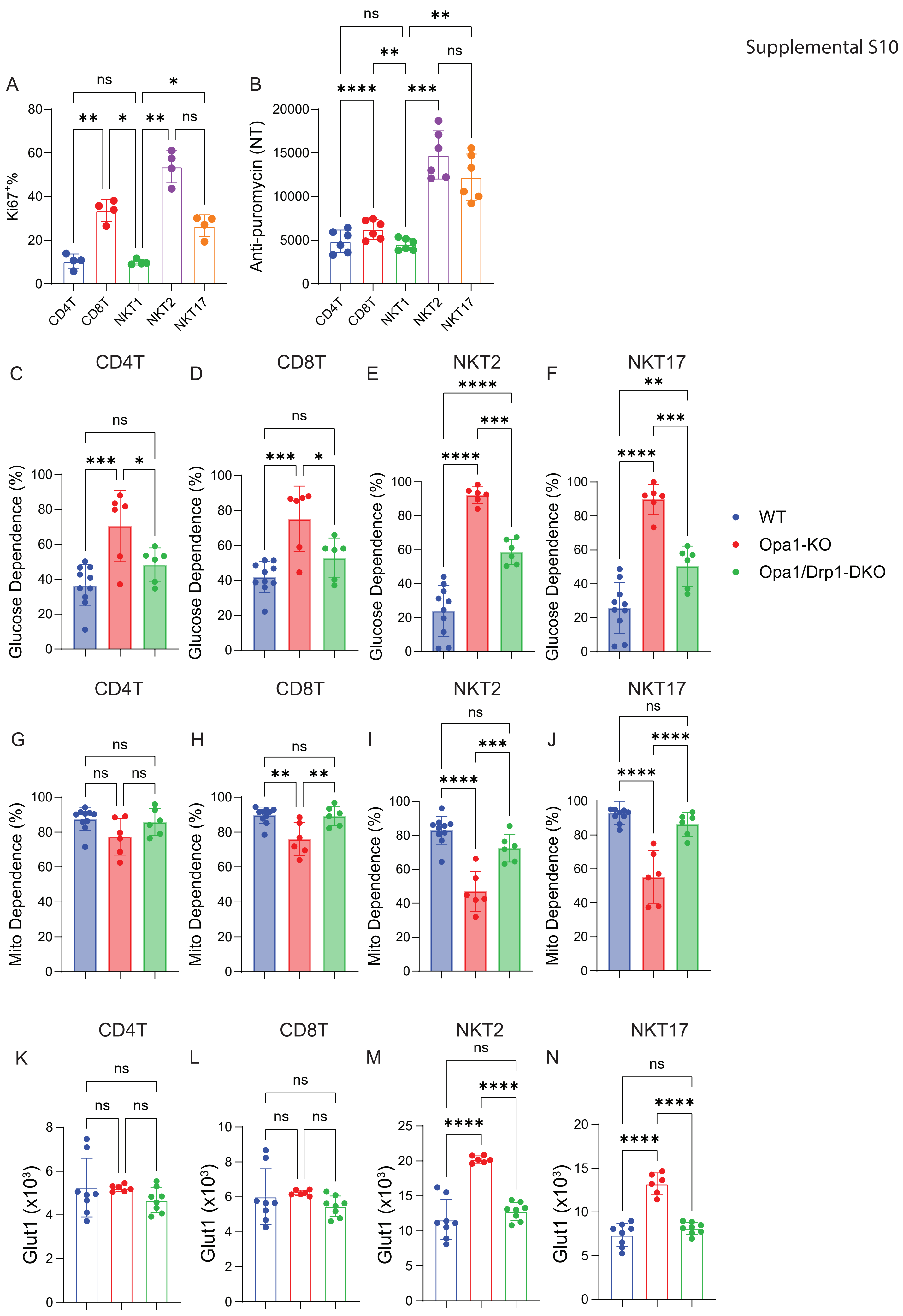

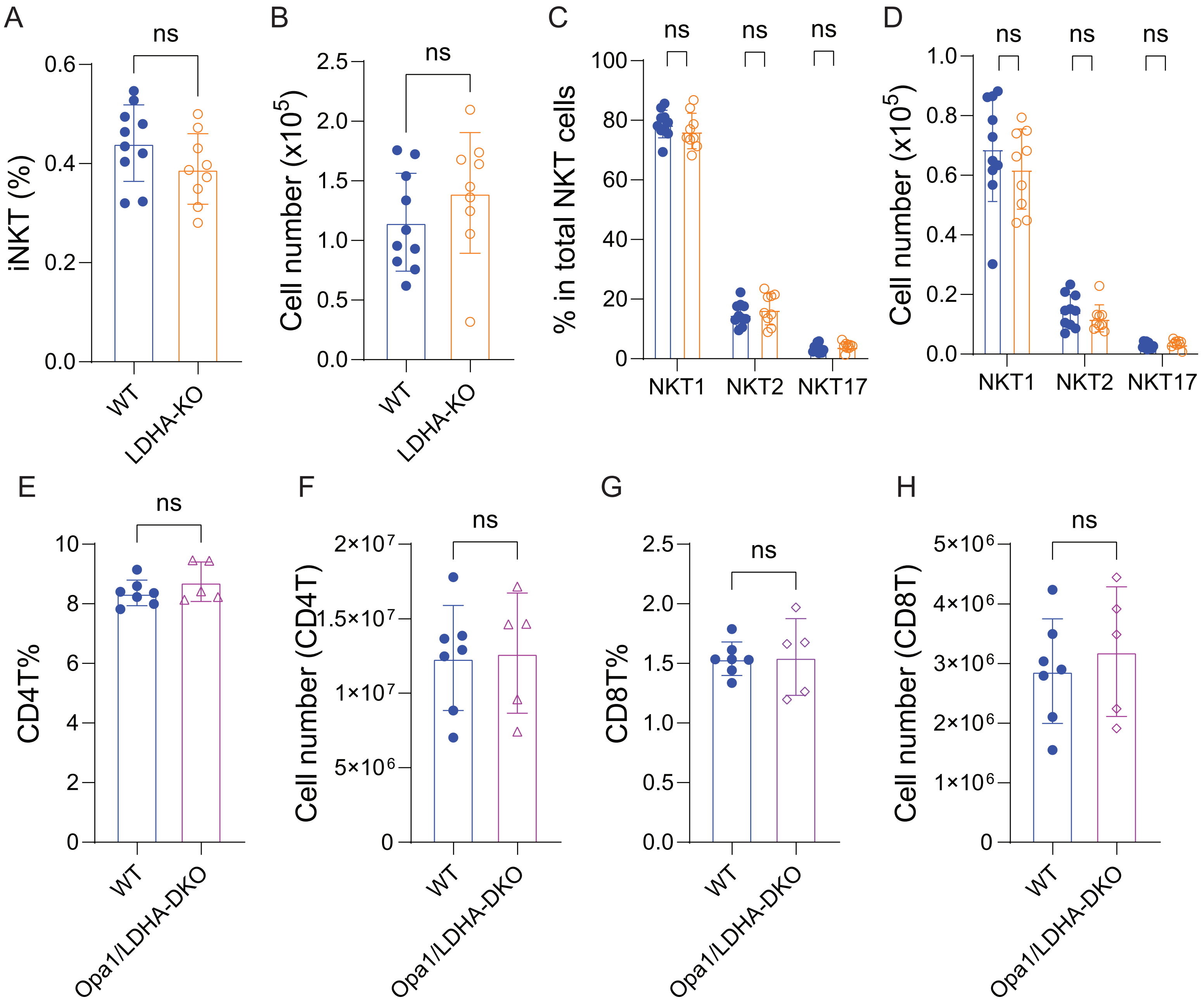

A
ns
0.6
0.4
0.2
0.0
iNKT (%)
WT
LDHA-KO
B
ns
2.5
2.0
1.5
1.0
0.5
0.0
Cell number (x10$^5$)
WT
LDHA-KO
C
ns
ns
ns
100
80
60
40
20
0
% in total NKT cells
NKT1
NKT2
NKT17
D
ns
ns
ns
1.0
0.8
0.6
0.4
0.2
0.0
Cell number (x10$^5$)
NKT1
NKT2
NKT17
E
ns
10
8
6
4
2
0
CD4T%
WT
Opa1/LDHA-DKO
F
ns
2×10$^7$
1.5×10$^7$
1×10$^7$
5×10$^6$
0
Cell number (CD4T)
WT
Opa1/LDHA-DKO
G
ns
2.5
2.0
1.5
1.0
0.5
0.0
CD8T%
WT
Opa1/LDHA-DKO
H
ns
5×10$^6$
4×10$^6$
3×10$^6$
2×10$^6$
1×10$^6$
0
Cell number (CD8T)
WT
Opa1/LDHA-DKO

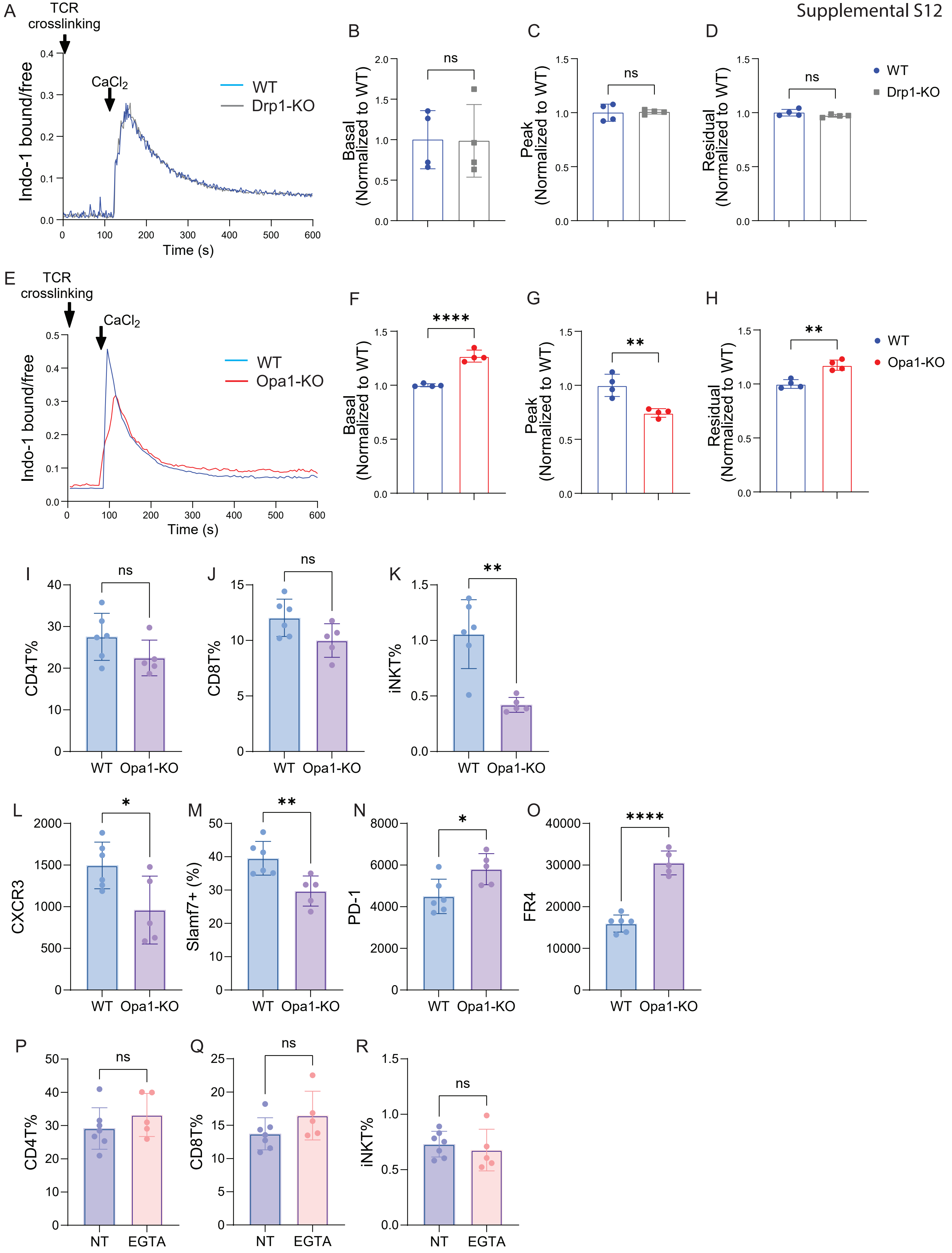
A
TCR crosslinking
CaCl2
WT
Drp1-KO
Indo-1 bound/free
Time (s)
B
Basal (Normalized to WT)
ns
C
Peak (Normalized to WT)
ns
D
Residual (Normalized to WT)
ns
WT
Drp1-KO
E
TCR crosslinking
CaCl2
WT
Opa1-KO
Indo-1 bound/free
Time (s)
F
Basal (Normalized to WT)
****
G
Peak (Normalized to WT)
**
H
Residual (Normalized to WT)
**
WT
Opa1-KO
I
CD4T%
ns
J
CD8T%
ns
K
iNKT%
**
WT Opa1-KO
L
CXCR3
*
M
Slamf7+ (%)
**
N
PD-1
*
O
FR4
****
WT Opa1-KO
P
CD4T%
ns
Q
CD8T%
ns
R
iNKT%
ns
NT EGTA

### Supplementary Table 1. Gene level membership of transcriptional modules

*Each row represents one gene–module membership. 'Detected in dataset' indicates presence in the analyzed scRNA-seq expression matrix.*

| Module ID | Gene | Detected in dataset | Provenance |
|---|---|---|---|
| mtDNA OXPHOS | mt-Co1 | Yes | mtDNA encoded OXPHOS genes detected in the scRNAseq dataset |
| mtDNA OXPHOS | mt-Cytb | Yes | mtDNA encoded OXPHOS genes detected in the scRNAseq dataset |
| mtDNA OXPHOS | mt-Nd5 | Yes | mtDNA encoded OXPHOS genes detected in the scRNAseq dataset |
| mtDNA OXPHOS | mt-Nd1 | Yes | mtDNA encoded OXPHOS genes detected in the scRNAseq dataset |
| mtDNA OXPHOS | mt-Nd2 | Yes | mtDNA encoded OXPHOS genes detected in the scRNAseq dataset |
| mtDNA OXPHOS | mt-Nd4 | Yes | mtDNA encoded OXPHOS genes detected in the scRNAseq dataset |
| Mitochondrial stress and proteostasis | Atf5 | Yes | Study-curated from bulk RNA-seq differences and literature-supported pathway genes (Quiros et al. J Cell Biol 2017; Munch & Harper, Nature 2016; Forsstrom et al. Cell Meta 2019); restricted to genes detected in the scRNAseq dataset |
| Mitochondrial stress and proteostasis | Txnrd2 | Yes | Study-curated from bulk RNA-seq differences and literature-supported pathway genes (Quiros et al. J Cell Biol 2017; Munch & Harper, Nature 2016; Forsstrom et al. Cell Meta 2019); restricted to genes detected in the scRNAseq dataset |
| Mitochondrial stress and proteostasis | Clpb | Yes | Study-curated from bulk RNA-seq differences and literature-supported pathway genes (Quiros et al. J Cell Biol 2017; Munch & Harper, Nature 2016; Forsstrom et al. Cell Meta 2019); restricted to genes detected in the scRNAseq dataset |
| Mitochondrial stress and proteostasis | Xpnpep3 | Yes | Study-curated from bulk RNA-seq differences and literature-supported pathway genes (Quiros et al. J Cell Biol 2017; Munch & Harper, Nature 2016; Forsstrom et al. Cell Meta 2019); restricted to genes detected in the scRNAseq dataset |
| Mitochondrial stress and proteostasis | Ppargc1b | Yes | Study-curated from bulk RNA-seq differences and literature-supported pathway genes (Quiros et al. J Cell Biol 2017; Munch & Harper, Nature 2016; Forsstrom et al. Cell Meta 2019); restricted to genes detected in the scRNAseq dataset |
| Mitochondrial stress and proteostasis | Mrps6 | Yes | Study-curated from bulk RNA-seq differences and literature-supported pathway genes (Quiros et al. J Cell Biol 2017; Munch & Harper, Nature 2016; Forsstrom et al. Cell Meta 2019); restricted to genes detected in the scRNAseq dataset |
| Mitochondrial stress and proteostasis | Mrps27 | Yes | Study-curated from bulk RNA-seq differences and literature-supported pathway genes (Quiros et al. J Cell Biol 2017; Munch & Harper, Nature 2016; Forsstrom et al. Cell Meta 2019); restricted to genes detected in the scRNAseq dataset |
| Mitochondrial stress and proteostasis | Elac2 | Yes | Study-curated from bulk RNA-seq differences and literature-supported pathway genes (Quiros et al. J Cell Biol 2017; Munch & Harper, Nature 2016; Forsstrom et al. Cell Meta 2019); restricted to genes detected in the scRNAseq dataset |
| Mitochondrial stress and proteostasis | Nsun4 | Yes | Study-curated from bulk RNA-seq differences and literature-supported pathway genes (Quiros et al. J Cell Biol 2017; Munch & Harper, Nature 2016; Forsstrom et al. Cell Meta 2019); restricted to genes detected in the scRNAseq dataset |
| Mitochondrial stress and proteostasis | Agk | Yes | Study-curated from bulk RNA-seq differences and literature-supported pathway genes (Quiros et al. J Cell Biol 2017; Munch & Harper, Nature 2016; Forsstrom et al. Cell Meta 2019); restricted to genes detected in the scRNAseq dataset |
| Mitochondrial stress and proteostasis | Samm50 | Yes | Study-curated from bulk RNA-seq differences and literature-supported pathway genes (Quiros et al. J Cell Biol 2017; Munch & Harper, Nature 2016; Forsstrom et al. Cell Meta 2019); restricted to genes detected in the scRNAseq dataset |
| Mitochondrial stress and proteostasis | Gfm1 | Yes | Study-curated from bulk RNA-seq differences and literature-supported pathway genes (Quiros et al. J Cell Biol 2017; Munch & Harper, Nature 2016; Forsstrom et al. Cell Meta 2019); restricted to genes detected in the scRNAseq dataset |
| Mitochondrial stress and proteostasis | Rexo2 | Yes | Study-curated from bulk RNA-seq differences and literature-supported pathway genes (Quiros et al. J Cell Biol 2017; Munch & Harper, Nature 2016; Forsstrom et al. Cell Meta 2019); restricted to genes detected in the scRNAseq dataset |
| Mitochondrial stress and proteostasis | Marchf5 | Yes | Study-curated from bulk RNA-seq differences and literature-supported pathway genes (Quiros et al. J Cell Biol 2017; Munch & Harper, Nature 2016; Forsstrom et al. Cell Meta 2019); restricted to genes detected in the scRNAseq dataset |
| Mitochondrial stress and proteostasis | Hspa9 | Yes | Study-curated from bulk RNA-seq differences and literature-supported pathway genes (Quiros et al. J Cell Biol 2017; Munch & Harper, Nature 2016; Forsstrom et al. Cell Meta 2019); restricted to genes detected in the scRNAseq dataset |
| Mitochondrial stress and proteostasis | Hspd1 | Yes | Study-curated from bulk RNA-seq differences and literature-supported pathway genes (Quiros et al. J Cell Biol 2017; Munch & Harper, Nature 2016; Forsstrom et al. Cell Meta 2019); restricted to genes detected in the scRNAseq dataset |
| Mitochondrial stress and proteostasis | Dnaja3 | Yes | Study-curated from bulk RNA-seq differences and literature-supported pathway genes (Quiros et al. J Cell Biol 2017; Munch & Harper, Nature 2016; Forsstrom et al. Cell Meta 2019); restricted to genes detected in the scRNAseq dataset |
| Mitochondrial stress and proteostasis | Dnajc11 | Yes | Study-curated from bulk RNA-seq differences and literature-supported pathway genes (Quiros et al. J Cell Biol 2017; Munch & Harper, Nature 2016; Forsstrom et al. Cell Meta 2019); restricted to genes detected in the scRNAseq dataset |
| Mitochondrial stress and proteostasis | Afg3l2 | Yes | Study-curated from bulk RNA-seq differences and literature-supported pathway genes (Quiros et al. J Cell Biol 2017; Munch & Harper, Nature 2016; Forsstrom et al. Cell Meta 2019); restricted to genes detected in the scRNAseq dataset |
| Mitochondrial stress and proteostasis | Pitrm1 | Yes | Study-curated from bulk RNA-seq differences and literature-supported pathway genes (Quiros et al. J Cell Biol 2017; Munch & Harper, Nature 2016; Forsstrom et al. Cell Meta 2019); restricted to genes detected in the scRNAseq dataset |
| Mitochondrial stress and proteostasis | Gfm2 | Yes | Study-curated from bulk RNA-seq differences and literature-supported pathway genes (Quiros et al. J Cell Biol 2017; Munch & Harper, Nature 2016; Forsstrom et al. Cell Meta 2019); restricted to genes detected in the scRNAseq dataset |
| Mitochondrial stress and proteostasis | Ptcd3 | Yes | Study-curated from bulk RNA-seq differences and literature-supported pathway genes (Quiros et al. J Cell Biol 2017; Munch & Harper, Nature 2016; Forsstrom et al. Cell Meta 2019); restricted to genes detected in the scRNAseq dataset |
| ISR/amino acid stress | Slc1a4 | Yes | Study-curated from bulk RNA-seq differences and literature-supported pathway genes (Quiros et al. J Cell Biol 2017; Munch & Harper, Nature 2016; Forsstrom et al. Cell Meta 2019); restricted to genes detected in the scRNAseq dataset |
| ISR/amino acid stress | Asns | Yes | Study-curated from bulk RNA-seq differences and literature-supported pathway genes (Quiros et al. J Cell Biol 2017; Munch & Harper, Nature 2016; Forsstrom et al. Cell Meta 2019); restricted to genes detected in the scRNAseq dataset |
| ISR/amino acid stress | Trib3 | Yes | Study-curated from bulk RNA-seq differences and literature-supported pathway genes (Quiros et al. J Cell Biol 2017; Munch & Harper, Nature 2016; Forsstrom et al. Cell Meta 2019); restricted to genes detected in the scRNAseq dataset |
| ISR/amino acid stress | Slc7a5 | Yes | Study-curated from bulk RNA-seq differences and literature-supported pathway genes (Quiros et al. J Cell Biol 2017; Munch & Harper, Nature 2016; Forsstrom et al. Cell Meta 2019); restricted to genes detected in the scRNAseq dataset |
| ISR/amino acid stress | Eif4ebp1 | Yes | Study-curated from bulk RNA-seq differences and literature-supported pathway genes (Quiros et al. J Cell Biol 2017; Munch & Harper, Nature 2016; Forsstrom et al. Cell Meta 2019); restricted to genes detected in the scRNAseq dataset |
| ISR/amino acid stress | Mthfd2 | Yes | Study-curated from bulk RNA-seq differences and literature-supported pathway genes (Quiros et al. J Cell Biol 2017; Munch & Harper, Nature 2016; Forsstrom et al. Cell Meta 2019); restricted to genes detected in the scRNAseq dataset |
| ISR/amino acid stress | Slc7a1 | Yes | Study-curated from bulk RNA-seq differences and literature-supported pathway genes (Quiros et al. J Cell Biol 2017; Munch & Harper, Nature 2016; Forsstrom et al. Cell Meta 2019); restricted to genes detected in the scRNAseq dataset |
| ISR/amino acid stress | Slc38a9 | Yes | Study-curated from bulk RNA-seq differences and literature-supported pathway genes (Quiros et al. J Cell Biol 2017; Munch & Harper, Nature 2016; Forsstrom et al. Cell Meta 2019); restricted to genes detected in the scRNAseq dataset |
| ISR/amino acid stress | Ggh | Yes | Study-curated from bulk RNA-seq differences and literature-supported pathway genes (Quiros et al. J Cell Biol 2017; Munch & Harper, Nature 2016; Forsstrom et al. Cell Meta 2019); restricted to genes detected in the scRNAseq dataset |
| ISR/amino acid stress | Ahcy | Yes | Study-curated from bulk RNA-seq differences and literature-supported pathway genes (Quiros et al. J Cell Biol 2017; Munch & Harper, Nature 2016; Forsstrom et al. Cell Meta 2019); restricted to genes detected in the scRNAseq dataset |
| ISR/amino acid stress | Shmt2 | Yes | Study-curated from bulk RNA-seq differences and literature-supported pathway genes (Quiros et al. J Cell Biol 2017; Munch & Harper, Nature 2016; Forsstrom et al. Cell Meta 2019); restricted to genes detected in the scRNAseq dataset |
| ISR/amino acid stress | Chdh | Yes | Study-curated from bulk RNA-seq differences and literature-supported pathway genes (Quiros et al. J Cell Biol 2017; Munch & Harper, Nature 2016; Forsstrom et al. Cell Meta 2019); restricted to genes detected in the scRNAseq dataset |
| ISR/amino acid stress | Slc3a2 | Yes | Study-curated from bulk RNA-seq differences and literature-supported pathway genes (Quiros et al. J Cell Biol 2017; Munch & Harper, Nature 2016; Forsstrom et al. Cell Meta 2019); restricted to genes detected in the scRNAseq dataset |
| ISR/amino acid stress | Asnsd1 | Yes | Study-curated from bulk RNA-seq differences and literature-supported pathway genes (Quiros et al. J Cell Biol 2017; Munch & Harper, Nature 2016; Forsstrom et al. Cell Meta 2019); restricted to genes detected in the scRNAseq dataset |
| ISR/amino acid stress | Mars1 | Yes | Study-curated from bulk RNA-seq differences and literature-supported pathway genes (Quiros et al. J Cell Biol 2017; Munch & Harper, Nature 2016; Forsstrom et al. Cell Meta 2019); restricted to genes detected in the scRNAseq dataset |
| ISR/amino acid stress | Aars | Yes | Study-curated from bulk RNA-seq differences and literature-supported pathway genes (Quiros et al. J Cell Biol 2017; Munch & Harper, Nature 2016; Forsstrom et al. Cell Meta 2019); restricted to genes detected in the scRNAseq dataset |
| ISR/amino acid stress | Sars | Yes | Study-curated from bulk RNA-seq differences and literature-supported pathway genes (Quiros et al. J Cell Biol 2017; Munch & Harper, Nature 2016; Forsstrom et al. Cell Meta 2019); restricted to genes detected in the scRNAseq dataset |
| ISR/amino acid stress | Iars | Yes | Study-curated from bulk RNA-seq differences and literature-supported pathway genes (Quiros et al. J Cell Biol 2017; Munch & Harper, Nature 2016; Forsstrom et al. Cell Meta 2019); restricted to genes detected in the scRNAseq dataset |
| ISR/amino acid stress | Nars | Yes | Study-curated from bulk RNA-seq differences and literature-supported pathway genes (Quiros et al. J Cell Biol 2017; Munch & Harper, Nature 2016; Forsstrom et al. Cell Meta 2019); restricted to genes detected in the scRNAseq dataset |
| ISR/amino acid stress | Cars | Yes | Study-curated from bulk RNA-seq differences and literature-supported pathway genes (Quiros et al. J Cell Biol 2017; Munch & Harper, Nature 2016; Forsstrom et al. Cell Meta 2019); restricted to genes detected in the scRNAseq dataset |
| ISR/amino acid stress | Dars | Yes | Study-curated from bulk RNA-seq differences and literature-supported pathway genes (Quiros et al. J Cell Biol 2017; Munch & Harper, Nature 2016; Forsstrom et al. Cell Meta 2019); restricted to genes detected in the scRNAseq dataset |
| ISR/amino acid stress | Farsb | Yes | Study-curated from bulk RNA-seq differences and literature-supported pathway genes (Quiros et al. J Cell Biol 2017; Munch & Harper, Nature 2016; Forsstrom et al. Cell Meta 2019); restricted to genes detected in the scRNAseq dataset |
| ISR/amino acid stress | Gars | Yes | Study-curated from bulk RNA-seq differences and literature-supported pathway genes (Quiros et al. J Cell Biol 2017; Munch & Harper, Nature 2016; Forsstrom et al. Cell Meta 2019); restricted to genes detected in the scRNAseq dataset |
| ISR/amino acid stress | Lars | Yes | Study-curated from bulk RNA-seq differences and literature-supported pathway genes (Quiros et al. J Cell Biol 2017; Munch & Harper, Nature 2016; Forsstrom et al. Cell Meta 2019); restricted to genes detected in the scRNAseq dataset |
| ISR/amino acid stress | Tars | Yes | Study-curated from bulk RNA-seq differences and literature-supported pathway genes (Quiros et al. J Cell Biol 2017; Munch & Harper, Nature 2016; Forsstrom et al. Cell Meta 2019); restricted to genes detected in the scRNAseq dataset |
| ISR/amino acid stress | Vars | Yes | Study-curated from bulk RNA-seq differences and literature-supported pathway genes (Quiros et al. J Cell Biol 2017; Munch & Harper, Nature 2016; Forsstrom et al. Cell Meta 2019); restricted to genes detected in the scRNAseq dataset |
| ISR/amino acid stress | Wars1 | Yes | Study-curated from bulk RNA-seq differences and literature-supported pathway genes (Quiros et al. J Cell Biol 2017; Munch & Harper, Nature 2016; Forsstrom et al. Cell Meta 2019); restricted to genes detected in the scRNAseq dataset |
| ISR/amino acid stress | Yars | Yes | Study-curated from bulk RNA-seq differences and literature-supported pathway genes (Quiros et al. J Cell Biol 2017; Munch & Harper, Nature 2016; Forsstrom et al. Cell Meta 2019); restricted to genes detected in the scRNAseq dataset |
| ISR/amino acid stress | Psat1 | Yes | Study-curated from bulk RNA-seq differences and literature-supported pathway genes (Quiros et al. J Cell Biol 2017; Munch & Harper, Nature 2016; Forsstrom et al. Cell Meta 2019); restricted to genes detected in the scRNAseq dataset |
| ISR/amino acid stress | Mthfd1l | Yes | Study-curated from bulk RNA-seq differences and literature-supported pathway genes (Quiros et al. J Cell Biol 2017; Munch & Harper, Nature 2016; Forsstrom et al. Cell Meta 2019); restricted to genes detected in the scRNAseq dataset |
| ISR/amino acid stress | Mthfs | Yes | Study-curated from bulk RNA-seq differences and literature-supported pathway genes (Quiros et al. J Cell Biol 2017; Munch & Harper, Nature 2016; Forsstrom et al. Cell Meta 2019); restricted to genes detected in the scRNAseq dataset |
| ISR/amino acid stress | Aldh18a1 | Yes | Study-curated from bulk RNA-seq differences and literature-supported pathway genes (Quiros et al. J Cell Biol 2017; Munch & Harper, Nature 2016; Forsstrom et al. Cell Meta 2019); restricted to genes detected in the scRNAseq dataset |
| ISR/amino acid stress | Bckdhb | Yes | Study-curated from bulk RNA-seq differences and literature-supported pathway genes (Quiros et al. J Cell Biol 2017; Munch & Harper, Nature 2016; Forsstrom et al. Cell Meta 2019); restricted to genes detected in the scRNAseq dataset |
| ISR/amino acid stress | Slc7a6 | Yes | Study-curated from bulk RNA-seq differences and literature-supported pathway genes (Quiros et al. J Cell Biol 2017; Munch & Harper, Nature 2016; Forsstrom et al. Cell Meta 2019); restricted to genes detected in the scRNAseq dataset |
| Terminal NKT1 | Ccl5 | Yes | Study-curated from published NKT1 signature genes (Engel et al., 2016; Krovi et al., 2020), restricted to genes detected in the scRNA-seq dataset. |
| Terminal NKT1 | Prf1 | Yes | Study-curated from published NKT1 signature genes (Engel et al., 2016; Krovi et al., 2020), restricted to genes detected in the scRNA-seq dataset. |
| Terminal NKT1 | Klrc1 | Yes | Study-curated from published NKT1 signature genes (Engel et al., 2016; Krovi et al., 2020), restricted to genes detected in the scRNA-seq dataset. |
| Terminal NKT1 | Klrb1c | Yes | Study-curated from published NKT1 signature genes (Engel et al., 2016; Krovi et al., 2020), restricted to genes detected in the scRNA-seq dataset. |
| Terminal NKT1 | Slamf7 | Yes | Study-curated from published NKT1 signature genes (Engel et al., 2016; Krovi et al., 2020), restricted to genes detected in the scRNA-seq dataset. |
| Terminal NKT1 | Il2rb | Yes | Study-curated from published NKT1 signature genes (Engel et al., 2016; Krovi et al., 2020), restricted to genes detected in the scRNA-seq dataset. |
| Terminal NKT1 | Ifng | Yes | Study-curated from published NKT1 signature genes (Engel et al., 2016; Krovi et al., 2020), restricted to genes detected in the scRNA-seq dataset. |
| Terminal NKT1 | Fasl | Yes | Study-curated from published NKT1 signature genes (Engel et al., 2016; Krovi et al., 2020), restricted to genes detected in the scRNA-seq dataset. |
| Terminal NKT1 | Runx3 | Yes | Study-curated from published NKT1 signature genes (Engel et al., 2016; Krovi et al., 2020), restricted to genes detected in the scRNA-seq dataset. |
| Terminal NKT1 | Il12rb2 | Yes | Study-curated from published NKT1 signature genes (Engel et al., 2016; Krovi et al., 2020), restricted to genes detected in the scRNA-seq dataset. |
| Terminal NKT1 | Il18rap | Yes | Study-curated from published NKT1 signature genes (Engel et al., 2016; Krovi et al., 2020), restricted to genes detected in the scRNA-seq dataset. |
| Terminal NKT1 | Ifngr1 | Yes | Study-curated from published NKT1 signature genes (Engel et al., 2016; Krovi et al., 2020), restricted to genes detected in the scRNA-seq dataset. |
| Terminal NKT1 | Itgae | Yes | Study-curated from published NKT1 signature genes (Engel et al., 2016; Krovi et al., 2020), restricted to genes detected in the scRNA-seq dataset. |
| Terminal NKT1 | Itga1 | Yes | Study-curated from published NKT1 signature genes (Engel et al., 2016; Krovi et al., 2020), restricted to genes detected in the scRNA-seq dataset. |
| Immature/NKT2-like | Tcf7 | Yes | Study-curated from published NKT1 signature genes (Engel et al., 2016; Krovi et al., 2020), restricted to genes detected in the scRNA-seq dataset. |
| Immature/NKT2-like | Lef1 | Yes | Study-curated from published NKT1 signature genes (Engel et al., 2016; Krovi et al., 2020), restricted to genes detected in the scRNA-seq dataset. |
| Immature/NKT2-like | Ccr9 | Yes | Study-curated from published NKT1 signature genes (Engel et al., 2016; Krovi et al., 2020), restricted to genes detected in the scRNA-seq dataset. |
| Immature/NKT2-like | Pak1 | Yes | Study-curated from published NKT1 signature genes (Engel et al., 2016; Krovi et al., 2020), restricted to genes detected in the scRNA-seq dataset. |
| Immature/NKT2-like | Ccr4 | Yes | Study-curated from published NKT1 signature genes (Engel et al., 2016; Krovi et al., 2020), restricted to genes detected in the scRNA-seq dataset. |
| Immature/NKT2-like | Sox4 | Yes | Study-curated from published NKT1 signature genes (Engel et al., 2016; Krovi et al., 2020), restricted to genes detected in the scRNA-seq dataset. |
| Immature/NKT2-like | Itpr1 | Yes | Study-curated from published NKT1 signature genes (Engel et al., 2016; Krovi et al., 2020), restricted to genes detected in the scRNA-seq dataset. |
| Immature/NKT2-like | Il17rb | Yes | Study-curated from published NKT1 signature genes (Engel et al., 2016; Krovi et al., 2020), restricted to genes detected in the scRNA-seq dataset. |
| Immature/NKT2-like | Zbtb16 | Yes | Study-curated from published NKT1 signature genes (Engel et al., 2016; Krovi et al., 2020), restricted to genes detected in the scRNA-seq dataset. |
| Immature/NKT2-like | Tox | Yes | Study-curated from published NKT1 signature genes (Engel et al., 2016; Krovi et al., 2020), restricted to genes detected in the scRNA-seq dataset. |
| Immature/NKT2-like | Il31ra | Yes | Study-curated from published NKT1 signature genes (Engel et al., 2016; Krovi et al., 2020), restricted to genes detected in the scRNA-seq dataset. |
| Immature/NKT2-like | Zbtb7b | Yes | Study-curated from published NKT1 signature genes (Engel et al., 2016; Krovi et al., 2020), restricted to genes detected in the scRNA-seq dataset. |
| Immature/NKT2-like | Rbpj | Yes | Study-curated from published NKT1 signature genes (Engel et al., 2016; Krovi et al., 2020), restricted to genes detected in the scRNA-seq dataset. |
| TCR/IEG (immediate early gene) | Nr4a1 | Yes | Study-curated genes representing the TCR-induced immediate-early response, including NR4A/AP-1 transcription factors and MAPK-responsive feedback genes. |
| TCR/IEG (immediate early gene) | Nr4a2 | Yes | Study-curated genes representing the TCR-induced immediate-early response, including NR4A/AP-1 transcription factors and MAPK-responsive feedback genes. |
| TCR/IEG (immediate early gene) | Nr4a3 | Yes | Study-curated genes representing the TCR-induced immediate-early response, including NR4A/AP-1 transcription factors and MAPK-responsive feedback genes. |
| TCR/IEG (immediate early gene) | Fos | Yes | Study-curated genes representing the TCR-induced immediate-early response, including NR4A/AP-1 transcription factors and MAPK-responsive feedback genes. |
| TCR/IEG (immediate early gene) | Fosb | Yes | Study-curated genes representing the TCR-induced immediate-early response, including NR4A/AP-1 transcription factors and MAPK-responsive feedback genes. |
| TCR/IEG (immediate early gene) | Fosl2 | Yes | Study-curated genes representing the TCR-induced immediate-early response, including NR4A/AP-1 transcription factors and MAPK-responsive feedback genes. |
| TCR/IEG (immediate early gene) | Jun | Yes | Study-curated genes representing the TCR-induced immediate-early response, including NR4A/AP-1 transcription factors and MAPK-responsive feedback genes. |
| TCR/IEG (immediate early gene) | Junb | Yes | Study-curated genes representing the TCR-induced immediate-early response, including NR4A/AP-1 transcription factors and MAPK-responsive feedback genes. |
| TCR/IEG (immediate early gene) | Jund | Yes | Study-curated genes representing the TCR-induced immediate-early response, including NR4A/AP-1 transcription factors and MAPK-responsive feedback genes. |
| TCR/IEG (immediate early gene) | Dusp1 | Yes | Study-curated genes representing the TCR-induced immediate-early response, including NR4A/AP-1 transcription factors and MAPK-responsive feedback genes. |
| TCR/IEG (immediate early gene) | Dusp5 | Yes | Study-curated genes representing the TCR-induced immediate-early response, including NR4A/AP-1 transcription factors and MAPK-responsive feedback genes. |
| TCR/IEG (immediate early gene) | Dusp10 | Yes | Study-curated genes representing the TCR-induced immediate-early response, including NR4A/AP-1 transcription factors and MAPK-responsive feedback genes. |
| TCR/IEG (immediate early gene) | Ier2 | Yes | Study-curated genes representing the TCR-induced immediate-early response, including NR4A/AP-1 transcription factors and MAPK-responsive feedback genes. |
| TCR/IEG (immediate early gene) | Ier5 | Yes | Study-curated genes representing the TCR-induced immediate-early response, including NR4A/AP-1 transcription factors and MAPK-responsive feedback genes. |
| NF-kB | Tnfaip3 | Yes | Study-curated genes representing the NF-κB transcriptional response, including inducible NF-κB targets/feedback regulators and NF-κB family members. |
| NF-kB | Nfkbia | Yes | Study-curated genes representing the NF-κB transcriptional response, including inducible NF-κB targets/feedback regulators and NF-κB family members. |
| NF-kB | Nfkbiz | Yes | Study-curated genes representing the NF-κB transcriptional response, including inducible NF-κB targets/feedback regulators and NF-κB family members. |
| NF-kB | Birc2 | Yes | Study-curated genes representing the NF-κB transcriptional response, including inducible NF-κB targets/feedback regulators and NF-κB family members. |
| NF-kB | Birc3 | Yes | Study-curated genes representing the NF-κB transcriptional response, including inducible NF-κB targets/feedback regulators and NF-κB family members. |
| NF-kB | Tnip1 | Yes | Study-curated genes representing the NF-κB transcriptional response, including inducible NF-κB targets/feedback regulators and NF-κB family members. |
| NF-kB | Rel | Yes | Study-curated genes representing the NF-κB transcriptional response, including inducible NF-κB targets/feedback regulators and NF-κB family members. |
| NF-kB | Relb | Yes | Study-curated genes representing the NF-κB transcriptional response, including inducible NF-κB targets/feedback regulators and NF-κB family members. |
| NF-kB | Nfkb2 | Yes | Study-curated genes representing the NF-κB transcriptional response, including inducible NF-κB targets/feedback regulators and NF-κB family members. |
| ISG (Interferon stimulated genes) | Isg15 | Yes | Canonical interferon-stimulated genes |
| ISG (Interferon stimulated genes) | Ifit1 | Yes | Canonical interferon-stimulated genes |
| ISG (Interferon stimulated genes) | Ifit3 | Yes | Canonical interferon-stimulated genes |
| ISG (Interferon stimulated genes) | Ifit3b | Yes | Canonical interferon-stimulated genes |
| ISG (Interferon stimulated genes) | Ifit1bl1 | Yes | Canonical interferon-stimulated genes |
| ISG (Interferon stimulated genes) | Irf7 | Yes | Canonical interferon-stimulated genes |
| ISG (Interferon stimulated genes) | Stat1 | Yes | Canonical interferon-stimulated genes |
| ISG (Interferon stimulated genes) | Stat2 | Yes | Canonical interferon-stimulated genes |
| ISG (Interferon stimulated genes) | Oasl2 | Yes | Canonical interferon-stimulated genes |
| ISG (Interferon stimulated genes) | Oasl1 | Yes | Canonical interferon-stimulated genes |
| ISG (Interferon stimulated genes) | Oas1a | Yes | Canonical interferon-stimulated genes |
| ISG (Interferon stimulated genes) | Oas2 | Yes | Canonical interferon-stimulated genes |
| ISG (Interferon stimulated genes) | Oas3 | Yes | Canonical interferon-stimulated genes |
| ISG (Interferon stimulated genes) | Rsad2 | Yes | Canonical interferon-stimulated genes |
| ISG (Interferon stimulated genes) | Usp18 | Yes | Canonical interferon-stimulated genes |
| ISG (Interferon stimulated genes) | Mx1 | Yes | Canonical interferon-stimulated genes |
| ISG (Interferon stimulated genes) | Mx2 | Yes | Canonical interferon-stimulated genes |
| ISG (Interferon stimulated genes) | Bst2 | Yes | Canonical interferon-stimulated genes |
| ISG (Interferon stimulated genes) | Ifi27l2a | Yes | Canonical interferon-stimulated genes |
| ISG (Interferon stimulated genes) | Ifi44 | Yes | Canonical interferon-stimulated genes |
| ISG (Interferon stimulated genes) | Ifi203 | Yes | Canonical interferon-stimulated genes |
| ISG (Interferon stimulated genes) | Ifi204 | Yes | Canonical interferon-stimulated genes |
| ISG (Interferon stimulated genes) | Ifi213 | Yes | Canonical interferon-stimulated genes |
| ISG (Interferon stimulated genes) | Iigp1 | Yes | Canonical interferon-stimulated genes |
| ISG (Interferon stimulated genes) | Igtp | Yes | Canonical interferon-stimulated genes |
| ISG (Interferon stimulated genes) | Gbp2 | Yes | Canonical interferon-stimulated genes |
| ISG (Interferon stimulated genes) | Gbp3 | Yes | Canonical interferon-stimulated genes |
| ISG (Interferon stimulated genes) | Gbp4 | Yes | Canonical interferon-stimulated genes |
| ISG (Interferon stimulated genes) | Gbp7 | Yes | Canonical interferon-stimulated genes |
| ISG (Interferon stimulated genes) | Zbp1 | Yes | Canonical interferon-stimulated genes |
| ISG (Interferon stimulated genes) | Xaf1 | Yes | Canonical interferon-stimulated genes |
| ISG (Interferon stimulated genes) | Cmpk2 | Yes | Canonical interferon-stimulated genes |
| ISG (Interferon stimulated genes) | Ifih1 | Yes | Canonical interferon-stimulated genes |
| ISG (Interferon stimulated genes) | Irf1 | Yes | Canonical interferon-stimulated genes |
| ISG (Interferon stimulated genes) | Irf9 | Yes | Canonical interferon-stimulated genes |
| ISG (Interferon stimulated genes) | Isg20 | Yes | Canonical interferon-stimulated genes |
| ISG (Interferon stimulated genes) | Ly6a | Yes | Canonical interferon-stimulated genes |
| ISG (Interferon stimulated genes) | Ly6c1 | Yes | Canonical interferon-stimulated genes |
| ISG (Interferon stimulated genes) | Ly6c2 | Yes | Canonical interferon-stimulated genes |
| ISG (Interferon stimulated genes) | Slfn1 | Yes | Canonical interferon-stimulated genes |
| ISG (Interferon stimulated genes) | Slfn5 | Yes | Canonical interferon-stimulated genes |
| ISG (Interferon stimulated genes) | Slfn8 | Yes | Canonical interferon-stimulated genes |
| ISG (Interferon stimulated genes) | Tap1 | Yes | Canonical interferon-stimulated genes |
| ISG (Interferon stimulated genes) | Tap2 | Yes | Canonical interferon-stimulated genes |
| ISG (Interferon stimulated genes) | Psmb8 | Yes | Canonical interferon-stimulated genes |
| ISG (Interferon stimulated genes) | Psmb9 | Yes | Canonical interferon-stimulated genes |
| ISG (Interferon stimulated genes) | B2m | Yes | Canonical interferon-stimulated genes |
| ISG (Interferon stimulated genes) | H2-K1 | Yes | Canonical interferon-stimulated genes |
| ISG (Interferon stimulated genes) | H2-D1 | Yes | Canonical interferon-stimulated genes |
| Proliferation | Kntc1 | Yes | Derived from the present scRNA-seq dataset (WT SC5 vs all other WT NKT1 subclusters). |
| Proliferation | Mcm10 | Yes | Derived from the present scRNA-seq dataset (WT SC5 vs all other WT NKT1 subclusters). |
| Proliferation | Pole | Yes | Derived from the present scRNA-seq dataset (WT SC5 vs all other WT NKT1 subclusters). |
| Proliferation | Uhrf1 | Yes | Derived from the present scRNA-seq dataset (WT SC5 vs all other WT NKT1 subclusters). |
| Proliferation | Rad51b | Yes | Derived from the present scRNA-seq dataset (WT SC5 vs all other WT NKT1 subclusters). |
| Proliferation | Dtl | Yes | Derived from the present scRNA-seq dataset (WT SC5 vs all other WT NKT1 subclusters). |

| | | | |
|---|---|---|---|
| Proliferation | Brca1 | Yes | Derived from the present scRNA-seq dataset (WT SC5 vs all other WT NKT1 subclusters). |
| Proliferation | Cit | Yes | Derived from the present scRNA-seq dataset (WT SC5 vs all other WT NKT1 subclusters). |
| Proliferation | Chaf1a | Yes | Derived from the present scRNA-seq dataset (WT SC5 vs all other WT NKT1 subclusters). |
| Proliferation | Hells | Yes | Derived from the present scRNA-seq dataset (WT SC5 vs all other WT NKT1 subclusters). |
| Proliferation | Lig1 | Yes | Derived from the present scRNA-seq dataset (WT SC5 vs all other WT NKT1 subclusters). |
| Proliferation | Top2a | Yes | Derived from the present scRNA-seq dataset (WT SC5 vs all other WT NKT1 subclusters). |
| Proliferation | Brip1 | Yes | Derived from the present scRNA-seq dataset (WT SC5 vs all other WT NKT1 subclusters). |
| Proliferation | Pola1 | Yes | Derived from the present scRNA-seq dataset (WT SC5 vs all other WT NKT1 subclusters). |
| Proliferation | Ncapg2 | Yes | Derived from the present scRNA-seq dataset (WT SC5 vs all other WT NKT1 subclusters). |
| Proliferation | Mcm3 | Yes | Derived from the present scRNA-seq dataset (WT SC5 vs all other WT NKT1 subclusters). |
| Proliferation | Stmn1 | Yes | Derived from the present scRNA-seq dataset (WT SC5 vs all other WT NKT1 subclusters). |
| Proliferation | Cdca7 | Yes | Derived from the present scRNA-seq dataset (WT SC5 vs all other WT NKT1 subclusters). |
| Proliferation | Brca2 | Yes | Derived from the present scRNA-seq dataset (WT SC5 vs all other WT NKT1 subclusters). |
| Proliferation | Mms22l | Yes | Derived from the present scRNA-seq dataset (WT SC5 vs all other WT NKT1 subclusters). |
| Proliferation | Diaph3 | Yes | Derived from the present scRNA-seq dataset (WT SC5 vs all other WT NKT1 subclusters). |
| Proliferation | Mcm5 | Yes | Derived from the present scRNA-seq dataset (WT SC5 vs all other WT NKT1 subclusters). |
| Proliferation | Mcm2 | Yes | Derived from the present scRNA-seq dataset (WT SC5 vs all other WT NKT1 subclusters). |
| Proliferation | Timeless | Yes | Derived from the present scRNA-seq dataset (WT SC5 vs all other WT NKT1 subclusters). |
| Proliferation | Zgrf1 | Yes | Derived from the present scRNA-seq dataset (WT SC5 vs all other WT NKT1 subclusters). |
| Proliferation | Mcm7 | Yes | Derived from the present scRNA-seq dataset (WT SC5 vs all other WT NKT1 subclusters). |
| Proliferation | Smc2 | Yes | Derived from the present scRNA-seq dataset (WT SC5 vs all other WT NKT1 subclusters). |
| Proliferation | Mcm4 | Yes | Derived from the present scRNA-seq dataset (WT SC5 vs all other WT NKT1 subclusters). |
| Proliferation | Cbx5 | Yes | Derived from the present scRNA-seq dataset (WT SC5 vs all other WT NKT1 subclusters). |
| Proliferation | Dnmt1 | Yes | Derived from the present scRNA-seq dataset (WT SC5 vs all other WT NKT1 subclusters). |
| Proliferation | Prim2 | Yes | Derived from the present scRNA-seq dataset (WT SC5 vs all other WT NKT1 subclusters). |
| Proliferation | Mcm6 | Yes | Derived from the present scRNA-seq dataset (WT SC5 vs all other WT NKT1 subclusters). |
| Proliferation | Cep128 | Yes | Derived from the present scRNA-seq dataset (WT SC5 vs all other WT NKT1 subclusters). |
| Proliferation | Blm | Yes | Derived from the present scRNA-seq dataset (WT SC5 vs all other WT NKT1 subclusters). |
| Proliferation | Dnajc9 | Yes | Derived from the present scRNA-seq dataset (WT SC5 vs all other WT NKT1 subclusters). |
| Proliferation | Tfdp1 | Yes | Derived from the present scRNA-seq dataset (WT SC5 vs all other WT NKT1 subclusters). |
| Proliferation | Nsd2 | Yes | Derived from the present scRNA-seq dataset (WT SC5 vs all other WT NKT1 subclusters). |
| Proliferation | Topbp1 | Yes | Derived from the present scRNA-seq dataset (WT SC5 vs all other WT NKT1 subclusters). |
| Proliferation | Ulbp1 | Yes | Derived from the present scRNA-seq dataset (WT SC5 vs all other WT NKT1 subclusters). |
| Proliferation | Tubb5 | Yes | Derived from the present scRNA-seq dataset (WT SC5 vs all other WT NKT1 subclusters). |
| Proliferation | Rrm1 | Yes | Derived from the present scRNA-seq dataset (WT SC5 vs all other WT NKT1 subclusters). |
| Proliferation | Ncapd3 | Yes | Derived from the present scRNA-seq dataset (WT SC5 vs all other WT NKT1 subclusters). |
| Proliferation | Atad2 | Yes | Derived from the present scRNA-seq dataset (WT SC5 vs all other WT NKT1 subclusters). |
| Proliferation | Nap1l1 | Yes | Derived from the present scRNA-seq dataset (WT SC5 vs all other WT NKT1 subclusters). |
| Proliferation | Usp37 | Yes | Derived from the present scRNA-seq dataset (WT SC5 vs all other WT NKT1 subclusters). |
| Proliferation | Psip1 | Yes | Derived from the present scRNA-seq dataset (WT SC5 vs all other WT NKT1 subclusters). |
| Proliferation | Fry | Yes | Derived from the present scRNA-seq dataset (WT SC5 vs all other WT NKT1 subclusters). |
| Proliferation | Ctdspl2 | Yes | Derived from the present scRNA-seq dataset (WT SC5 vs all other WT NKT1 subclusters). |
| Proliferation | Rbbp8 | Yes | Derived from the present scRNA-seq dataset (WT SC5 vs all other WT NKT1 subclusters). |
